# SUMO2 Protects Against Tau-induced Synaptic and Cognitive Dysfunction

**DOI:** 10.1101/2022.11.11.516192

**Authors:** Franca Orsini, Elentina Argyrousi, Elena Restelli, Lenzie K. Ford, Hironori Takamura, Shinsuke Matsuzaki, Lorena Zentilin, Rosaria Pascente, Nicholas M Kanaan, Rajesh Soni, Taiichi Katayama, Roberto Chiesa, Gianluigi Forloni, Kenneth S. Kosik, Eric R. Kandel, Paul E. Fraser, Ottavio Arancio, Luana Fioriti

## Abstract

Abnormal intracellular accumulation of Tau aggregates is a hallmark of Alzheimer’s disease (AD) and other Tauopathies, such as Frontotemporal dementia (FTD), which can be caused by mutations of Tau. Mutated and pathological Tau can undergo a range of post-translational modifications (PTMs) that might trigger or modulate disease pathology. Recent studies indicate that modification of wild type Tau by <u>S</u>mall <u>u</u>biquitin-like <u>m</u>odifier SUMO isoform 1 (SUMO1) controls Tau hyperphosphorylation and aggregation, suggesting that SUMOylation acts as a central regulator of Tau’s biochemical properties. Besides SUMO1, Tau is modified by SUMO2/3, however the consequences of this modification have not been investigated. Here, using viral approaches on primary hippocampal neurons, transgenic mice expressing mutant Tau and SUMO2, and iPSC-derived neurons from FTD patients, we evaluated whether SUMO2/3 conjugation modifies the neurodegenerative disease pathology associated with the aggregation-prone mutant Tau P301L, P301S, and R406W variants. We found that mutant forms of Tau are targets of SUMO2/3, and SUMO2/3 conjugation is neuroprotective. Importantly, expression of mutant Tau is accompanied by a significant reduction of SUMO2/3 conjugation levels, and restoring levels of SUMO2 reduces mutant Tau aggregation and phosphorylation in all model systems Furthermore, overexpression of SUMO2 restores levels of pre- and post-synaptic markers, associated with a complete rescue of the LTP and memory deficits in transgenic mice expressing mutant Tau. These findings bring to light the potential therapeutic implication of manipulating SUMO conjugation to detoxify Tau through PTM-based approaches.

## INTRODUCTION

The microtubule-associated protein Tau is highly expressed in the central nervous system and is linked to the pathology of several neurodegenerative diseases, jointly termed Tauopathies. Although Tau was found to participate in regulating microtubules dynamics and axonal transport in neurons ^1^, the full physiological function of Tau remains unknown. In adult neurons, six isoforms of Tau exist in a homeostatic balance^2^. Mutations of the Tau encoding gene (*MAPT*), abnormal isoforms expression, abnormal phosphorylation and/or conformational changes of the Tau protein can disturb its physiological balance, leading to Tau pathologies ^3^. Tauopathies present diverse clinical phenotypes ranging from dementia and psychiatric disorders to movement disorders^4^. Among the most common forms of Tauopathies are Frontotemporal dementia (FTD) and Alzheimer’s disease (AD). They share both clinical and neuropathological features, such as neurofibrillary tangles, which consist of aggregated and abnormally phosphorylated Tau deposits ^5 6 7^. Several pathogenic mutations in the Tau gene were identified in families with FTD and parkinsonism linked to chromosome 17 (FTDP-17) ^3^. Among the well-studied mutations are P301S, P301L, and R406W, for which animal models of the disease exist^8^. Some of these mutations promote toxicity by enhancing Tau aggregation, interfering with Tau’s microtubule-dependent and independent functions ^9^, and cause synaptic loss. However, the molecular mechanisms behind these phenomena are still poorly understood.

Tau is subject to several post-translational modifications (PTM)^10^, including phosphorylation, truncation, acetylation, N-glycosylation, nitrogenation, and ubiquitination. While phosphorylation of Tau are widely investigated and alterations have been associated with reduced binding to microtubules ^11^ and mis-sorting of Tau to dendrites and dendritic spines ^12^, much less is known about the functional consequences of other PTMs, such as SUMOylation. SUMOylation is a covalent and reversible attachment of an 11 kDa SUMO (Small Ubiquitin-like MOdifier) protein to a lysine residue ^13 14^. The covalent attachment of the SUMO protein represents a critically important control process in all eukaryotic cells, as it acts as a biochemical switch to regulate the function of hundreds of proteins in many different pathways. Although the functional consequences of SUMOylation are manyfold, the underlying principle is that it alters the inter- and/or intra-molecular interactions of substrate proteins and thus modulates their activity, stability, and subcellular localization ^14^.

In the brain of vertebrates, three main SUMO isoforms (SUMO1-3) are characterized. While SUMO-2 and SUMO-3 differ by only three N-terminal amino acids and are often referred to as SUMO2/3, they only share ~ 50% of their amino acid sequences with SUMO1 ^15^. Moreover, significant differences are observed in the functions of SUMO1 and SUMO2/3. Under resting conditions, for example, very little unconjugated SUMO1 is present, whereas a large free pool of SUMO-2/3 is available ^16^, which becomes conjugated to target proteins in response to cellular stress ^17 18^. This increase in SUMO2/3 protein conjugation confers resistance to stress. In particular neurons depend on SUMO2/3 conjugation to withstand stress conditions, such as ischemia ^19 20^ and amyloid-induced toxicity ^21 22^.

SUMOylation has been implicated in the pathogenesis of several neurodegenerative disorders including Huntington’s ^23^ and other polyglutamine diseases ^24 25^ Parkinson’s disease ^26 27 28^ and AD ^22 29 30 31 32^. Post-mortem tissue samples from the hippocampal formation of AD patients exhibited significantly less high molecular weight SUMO2/3 conjugates compared to controls ^33^. This dysregulation was specific for SUMO2/3, as levels of SUMO1 conjugates were not significantly altered ^33^.

Despite the evidence of dysregulated SUMOylation in secondary Tauopathies, such as AD, the contribution of SUMOylation to the pathogenesis of primary Tauopathies, such as FTD, remains unknown. Here we investigated whether global SUMOylation is affected in two animal models of FTD, the JNPL3 ^34^ and PS19 mice ^35^, as well as in human iPSC-derived neurons expressing mutant forms of Tau (R406W and P301L). In all systems, we found that SUMO2/3 conjugation is reduced.

SUMO1 promotes phosphorylation and aggregation of wild-type Tau ^36^ but the effects of SUMO2/3 conjugation are unknown. We previously found that SUMO1 and SUMO2 act differently on the prion-like protein CPEB3 ^37 38^, with SUMO1 promoting and SUMO2 reducing CPEB3 oligomerization^39^. Here, we present data obtained *in vitro* and *in vivo* showing that at least some forms of mutant Tau (P301L/S) are *bona fide* targets of both SUMO1 and SUMO2. We also show that SUMO2 reduced mutant Tau aggregation and protected against mutant Tau (P301L/S)-induced synaptic toxicity. Finally, we find that overexpression of SUMO2 completely rescued the learning deficits and restored synaptic plasticity in mice expressing mutant Tau.

Overall, our data implicate dysregulated SUMOylation as a novel factor in FTD pathology, and provide compelling evidence that SUMO2 protects neurons from Tau toxicity.

## Results

### Global SUMO conjugation is altered in animal models of FTD

Global levels of SUMOylated proteins are increased in a number of neurodegenerative diseases; however, no data are currently available on global SUMOylation in FTD. Here we compared the profile of SUMO conjugation in two animal models of the disease, the transgenic mice PS19 ^34^ and JNPL3 ^35^, which express mutant forms of Tau associated with primary Tauopathies, namely P301S and P301L, respectively. We quantified SUMOylated proteins in cortical extracts of 6-9 month old mice and littermate controls. We found increased levels of SUMO1 conjugated proteins and decreased levels of SUMO2 in the brains of Tg mice expressing mutant Tau (Fig. 1A and Supplemental Fig. 1). The imbalance of SUMO1 and SUMO2 conjugation might affect multiple cellular pathways important for the onset of the disease ^13,40^. We used SUMO-immunoprecipitation (IP, using a recently developed protocol ^41^) to identify key SUMOylated proteins (Fig. 1B) in an unbiased fashion through mass spectrometry. Interestingly, Tau was one of the most abundantly modified proteins, suggesting that mutant Tau might be a direct target of SUMOylation. Similar findings were obtained on transgenic mice expressing SUMO1 ^42^, where wild type Tau was found to be a prime target of SUMO1 conjugation.

**Fig. 1.**
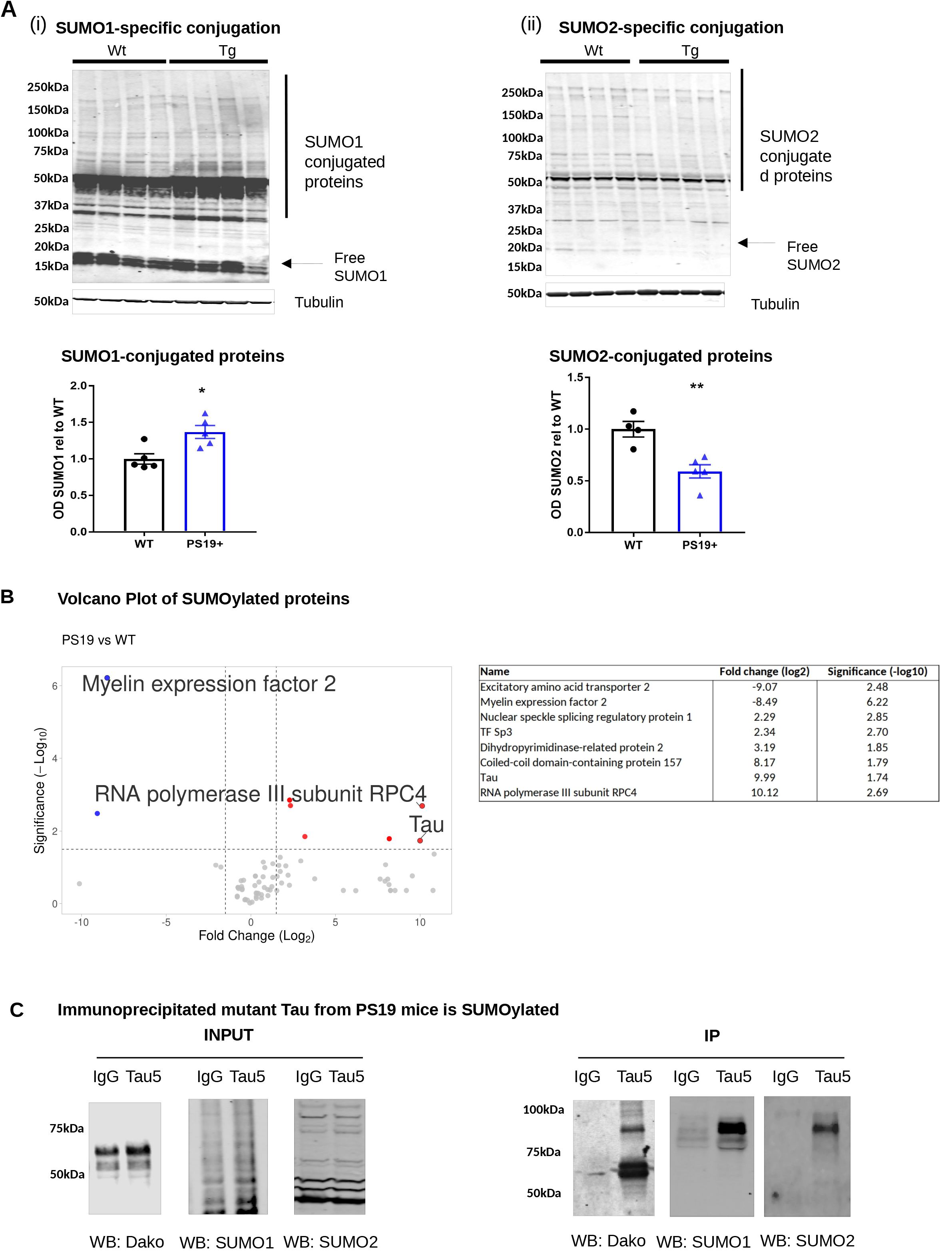
Global SUMO conjugation is altered in models of FTD. **A)** Representative western blot of global SUMO1 (i) and SUMO2 (ii) conjugation in 7-9 months old PS19 (Tg) and Wt (Non-Tg littermate) mice. Total cortical extracts were probed with SUMO1 or SUMO2 specific antibodies in 4-5 animals per genotype. PS19 show significant increase of SUMO1 conjugation and reduction of SUMO2 conjugation compared to their respective controls, * p<0.05 and ** p<0.01, unpaired t-test, data are mean ± SEM. Tubulin was used as loading control. **B)** Volcano plot of differently SUMOylated proteins in cortical extracts of Wt and PS19 mice processed with SUMO remnant immuno profiling method ^41^. Red dots indicate increased SUMOylated proteins while the blue dots correspond to decreased sumoylated protein in PS19 vs Wt, as listed in the Table. **C)** Immunoprecipitation experiments of mutant Tau using total tau antibody (tau5) from PS19 cortices followed by western blot with anti tau (Dako antibody) and SUMO1 and SUMO2 (Enzo) identify SUMO1- and SUMO2 conjugated Tau.

### Mutant P301S Tau is SUMOylated in brain extracts of FTD mice

SUMOylation of wild type Tau was reported by several groups *in vitro* ^36 43^ and recently also *in vivo* ^44^; however, there are no data on SUMOylation of mutant Tau or the consequences of mutant Tau SUMOylation. To confirm our mass spec results on mutant Tau SUMOylation, we performed an *in vitro* reconstituted SUMOylation assay using purified recombinant proteins. We used recombinant human Tau (hTau441) and mutant Tau (P301L and P301S) purified from bacteria ^45^ and found that both mutant forms of Tau were SUMOylated by SUMO1 and SUMO2 in an ATP-dependent manner, similar to wild-type Tau (Supplemental Fig. 1B).

To further confirm that SUMOylation of mutant Tau also occurs *in vivo,* we performed IP experiments using hippocampal tissue of transgenic mice expressing P301S Tau. To this end, 8-9 months old mice were sacrificed and their hippocampi were isolated. Protein extracts were next subjected to IP using a total Tau antibody and the eluted samples were probed with SUMO1 and SUMO2-specific antibodies. We found that a band compatible with SUMOylated Tau was present (Fig. 1C). Together, these results indicate that mutant Tau is a *bona fide* target of the SUMOylation machinery.

### SUMO2 levels modulate aggregation of mutant Tau in HT22 cells

Next, we sought to determine the effects of SUMO conjugation on mutant forms of Tau. SUMO1 was reported to increase aggregation and toxicity of wild type Tau ^36^. To explore the role of SUMOylation in modulating aggregation and toxicity of mutant Tau, we co-expressed mutant TauP301L-HA (mut-Tau) with SUMO1-GFP or SUMO2-GFP in HT22 cells (Fig. 2A) of neuronal origin ^46 47^.

**Fig. 2.**
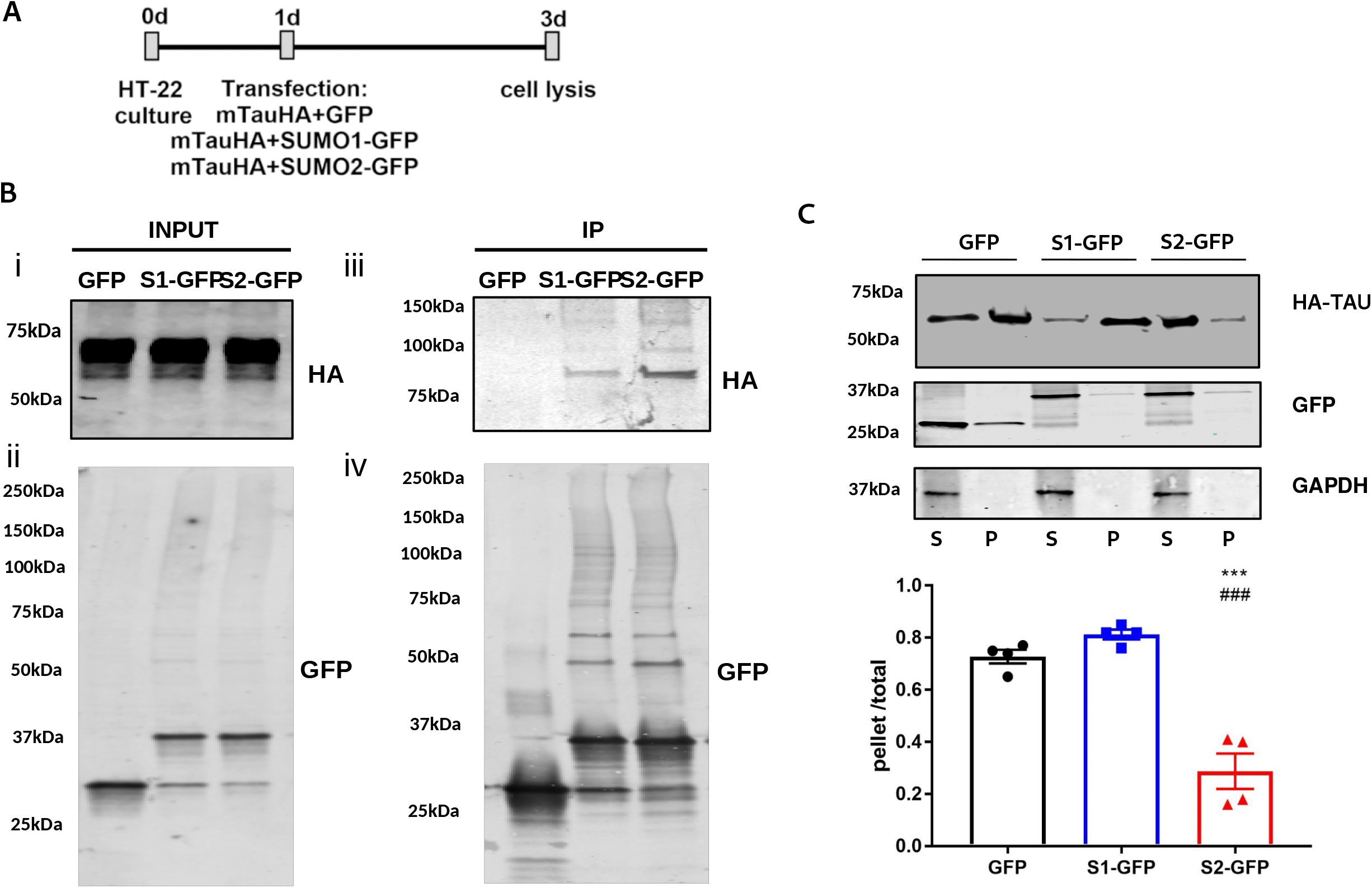
SUMO2 reduces the aggregation of TauP301L in HT22 neuronal cells. **A)** Experimental design: HT22 cells were transfected for 48h with TauP301L-HA in combination with GFP or SUMO1-GFP (S1-GFP) and SUMO2-GFP (S2-GFP). **B)** Total protein extracts demonstrated equal expression of the TauP301L-HA protein (i) and GFP (ii) in the different conditions SUMOylated proteins were immunoprecipitated using GFP and probed for HA (iii) and GFP (iv). **C)** Representative western blot showing detergent insolubility of TauP301L-HA detected with HA antibody. Membranes were also probed for GFP to check for S1- and S2-GFP expression, and GAPDH, as a control for ultracentrifugation procedure. Soluble and insoluble fractions are labeled as ‘S’ (sup) and ‘P’ (pellet) respectively. The graph shows the quantification of 4 independent experiments. Data are reported as mean± SEM., one way ANOVA with Tukey’s post hoc test, ***p<0.0001 (S2-GFP vs GFP and ###p<0.0001 S2-GFP vs S1-GFP).

We performed IP experiments and confirmed that mut-Tau is a target of both SUMO1 and SUMO2 in HT22 cells (Fig. 2B). To assess whether different SUMO moieties have different effects on mutant Tau aggregation, we subjected lysates from cells overexpressing mut-Tau with GFP or mut-Tau together with SUMO1-GFP or SUMO2-GFP to a detergent insolubility assay, a technique that measures the degree of solubility of proteins in non-ionic detergents ^48^. Similar to other amyloid and amyloid-like proteins ^37^, pathological Tau is partially insoluble in these buffers ^36^ when expressed at very high levels. We found that overexpression of SUMO2 decreases the insoluble and aggregated fraction of mut-Tau (Fig. 2C). In contrast, SUMO1 failed to reduce the insoluble fraction of mut-Tau. These data suggest that different SUMO isoforms exert different effects on Tau aggregation.

One of the caveats of overexpressing of SUMO variants is the broad effect, direct or indirect, they can have on hundreds of proteins. To establish whether the reduced aggregation of mutant Tau is due to a direct effect of SUMO2, we repeated our experiments using a form of mut-Tau that cannot be SUMOylated, the K340R mut-Tau ^36 43^. We found that preventing SUMOylation at K340 blocked the changes in mut-Tau aggregation induced by overexpression of SUMO2 (Supplemental Fig. 2).

Together our findings suggest that SUMO2 can directly prevent aggregation of mut-Tau, and that K340 is a critical modification site for this effect. Based on these results, we formulated the hypothesis that SUMO2 conjugation may ameliorate Tau-induced toxicity in neurons.

### SUMO2 decreases the aggregation and phosphorylation of TauP301L in hippocampal primary neurons *in vitro*

To test the hypothesis that SUMO2 might reduce the aggregation of mutant Tau in neurons, we cultured primary hippocampal neurons and overexpressed mut-Tau with a C-terminal HA tag together with GFP as control, or together with SUMO2-RFP, using a viral approach (Fig. 3A). Similar to what we found in HT22 cells, SUMO2 overexpression reduced mut-Tau aggregation (Fig. 3B), as shown by detergent insolubility assay. We confirmed these findings using an additional technique called SDD-AGE, semi denaturing agarose gel electrophoresis^49^, and found that the high molecular weight assemblies of mut-Tau are reduced in the presence of SUMO2 (Fig. 3C).

**Fig. 3.**
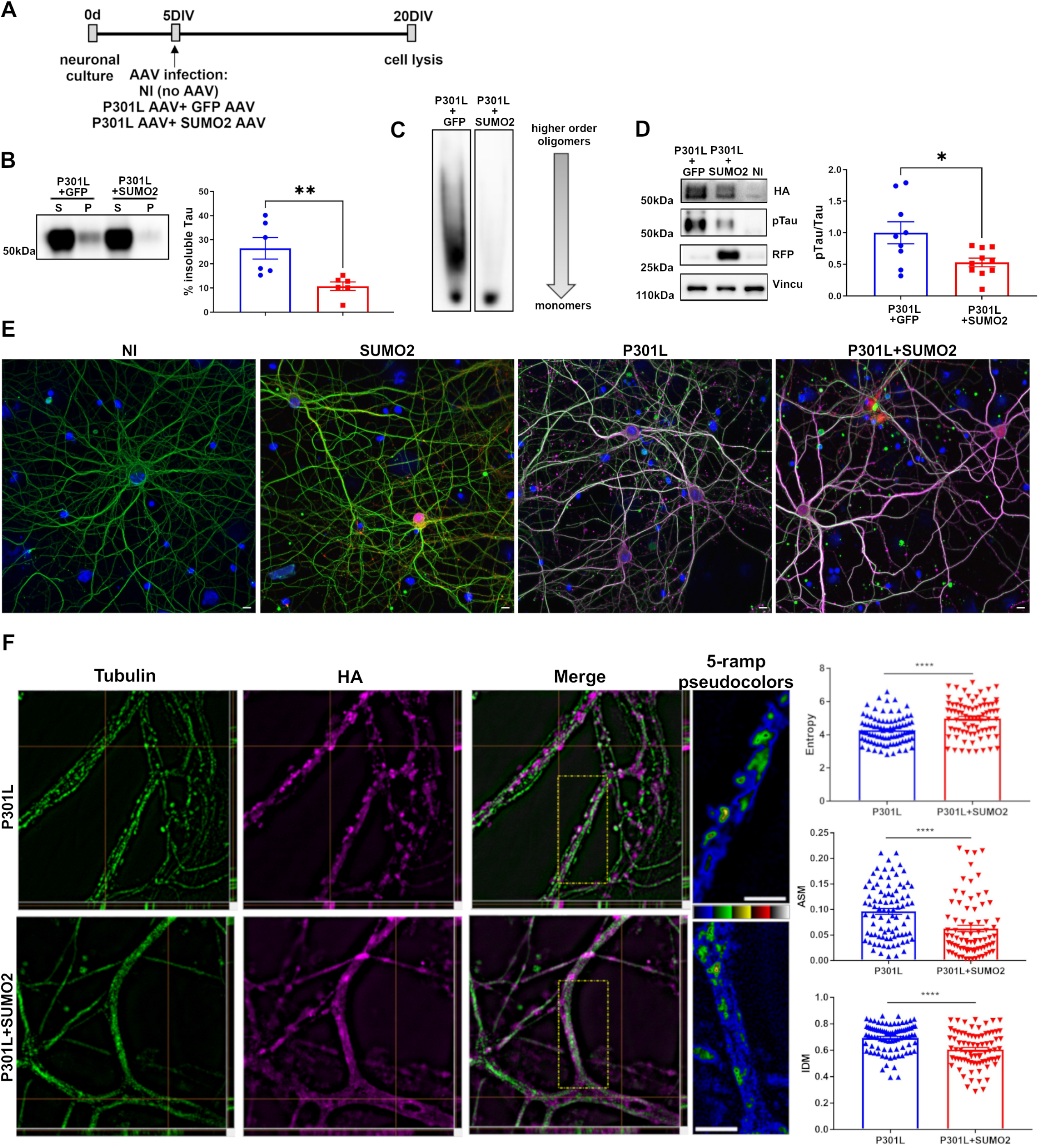
SUMO2 reduces the aggregation and phosphorylation of TauP301L in cultured primary hippocampal neurons. **A)** Experimental design: primary hippocampal neurons (HP) at DIV 5 were infected with AAV transducing 2N4R TauP301L-HA tag in combination with GFP AAV (to ensure equal viral load) or SUMO2 AAV and then lysed at day 20. **B)** Representative western blot probed with HA antibody showing detergent insolubility of TauP301L protein from neurons transduced with TauP301L in combination with GFP AAV or SUMO2 AAV. Soluble and pellet fractions are labeled as ‘S’ and ‘P’ respectively. The graph shows the quantification of 6 samples from 4 independent experiments. Data are the mean± SEM., Unpaired t-test, **p<0,01. **C)** Representative semi-denaturing detergent agarose gel of cell lysates from neurons transduced with TauP301L in combination with GFP AAV or SUMO2 AAV. Agarose gel probed with Tau-5A6 antibody shows a reduction of higher order oligomers in P301L+SUMO2 AAV compared P301L+GFP AAV transduced cells. **D)** Representative western blot probed with anti-phosphoTau (198/199/202/205) antibody showing significant reduction of hyperphosphorylation of Tau in neurons transduced with TauP301L in combination with SUMO2 AAV. Blots were probed with an anti-vinculin antibody as loading control. The graph shows the quantification of 9 samples from 4 independent experiments. Data are the mean± SEM., unpaired t-test,**p<0.01. **E)** representative images of hp neurons non infected (NI) SUMO2-RFP, P301L-HA and P301L-Ha + SUMO2-RFP infected. Tubulin staining is shown in green, SUMO2 in red and HA staining in magenta. Blue for DAPI. **F)** Representative super-resolution images of dendrites from transduced neurons show clustered HA signals (magenta) close to tubulin (green) in P301L AAV, and a more scattered HA signal in P301L AAV+ SUMO2 AAV. Right panels show magnification of one branch per image in ‘5 ramps’ modality (scale bar=2.5 um). Gray-level co-occurrence matrix (GLCM) was used for the quantification of HA signal distribution in dendrites. GLCM analysis showed higher entropy and lower angular second moment (ASM) and inverse difference moment (IDM) in P301L +SUMO2 AAVs neurons compared to P301L AAV alone, indicating lower pixel homogeneity corresponding to a reduction of clustered pixels. The graph shows the quantification of 88-78 branches for P301L and P301L+SUMO2 respectively from two independent experiments. Data are the mean± SEM, Mann-Whitney test,****p<0.0001.

Abnormal phosphorylation of Tau at specific residues is reported in patients affected by AD and FTD ^50^, as well as in several disease models ^51^. Some forms of phosphorylated Tau display lower affinity for microtubules and forms increased aggregation potential ^11^, suggesting a possible link between the degree of phosphorylation and formation of aggregates ^52^, which are toxic to neurons ^53^. Given the reduced propensity of mutant Tau to form aggregates when modified by SUMO2, we quantified the amount of phosphorylated Tau at residues 198, 199, 202 and 205 ^54^. We found a significant reduction of phosphorylated mut-Tau at these sites when SUMO2 was overexpressed (Fig. 3D). These results prompted us to assess the distribution of mutant Tau in our neuronal cultures. To this end we used AAV to transduce neurons with TauP301L-HA or SUMO2-RFP alone or in combination. After 15 days we stained cells with tubulin and tested the expression of mTau through HA staining. As expected, no HA signal was detected on non-infected (NI) and SUMO2-RFP groups, while a robust signal was observed in P301L and P301L+SUMO2 groups (Fig. 3E). In these 2 groups, we next took advantage of structure illuminated microscopy (SIM) to analyze mut-Tau distribution in hippocampal neurons. We found that mutant Tau appeared both in a diffused and clustered pattern (Fig. 3E). When SUMO2 was co-expressed, we found that mutant Tau had a more diffused signal. We quantified the degree of diffuse signal by performing image texture analysis, using the gray level co-occurrence matrix (GLCM) method. This allowed us to measure the entropy, angular second moment (ASM) and inverse difference moment (IDM) of the mutant Tau signal in neuronal processes ^55^. We found that overexpression of SUMO2 increased the entropy in cells expressing mut-Tau, compared to cells expressing mut-Tau alone (Fig. 3F). ASM is a measure of textural uniformity of an image. We found that ASM is lower when TauP301L is co-expressed with SUMO2. IDM measures image homogeneity. Again, we found that IDM was higher in cells expressing mut-Tau with SUMO2, indicating that the distribution of mutant Tau is more homogeneous and diffuse in the presence of SUMO2. As control, we performed the same analyses on endogenous wild type Tau using the antibody T49 and found no changes in distribution of endogenous Tau signal (Supplemental Fig. 3) when SUMO2 was overexpressed.

Next, we assessed the co-localization of mutant Tau (HA staining) with microtubules (tubulin staining). We calculated the Pearson correlation coefficient (PCC) and found a value of 0.7 between TauP301L (HA signal) and tubulin in cells expressing mut-Tau alone. This value significantly increased to 0.77 (p<0.05) in case of co-expression with SUMO2 (Supplemental Fig. 4). These data suggest that in the presence of SUMO2 mutant Tau is less clustered and more associated with microtubules.

### SUMO2 reduces the amount of TauP301L in Post-synaptic densities of primary hippocampal neurons *in vitro*

Tau is normally distributed throughout the neuron (i.e. soma, dendrites, nucleus and axon) ^55^, however, pathological species of tau can accumulate in dendritic spines and are thought to cause synaptic dysfunction 35 ^56 57^. –Given that increased SUMO2 levels reduced the aggregation of mut-Tau, we next sought to determine whether increased SUMO2 also affected the degree to which mut-Tau accumulates at the synapse. We quantified on SIM images the degree of colocalization of TauP301L with PSD95, a marker of spines, or synaptophysin, a marker of pre-synaptic sites, in cultured primary hippocampal neurons infected with mut-Tau alone, or together with SUMO2-RFP. In the presence of SUMO2, the co-localization of mut-Tau with PSD95 was significantly reduced, suggesting that mut-Tau is less present in dendritic spines, compared to expression of mut-Tau alone (Fig. 4A). We also found that the degree of colocalization between mut-Tau and synaptophysin did not change, suggesting the SUMO2 did not affect presynaptic localization of mut-Tau.

**Fig. 4.**
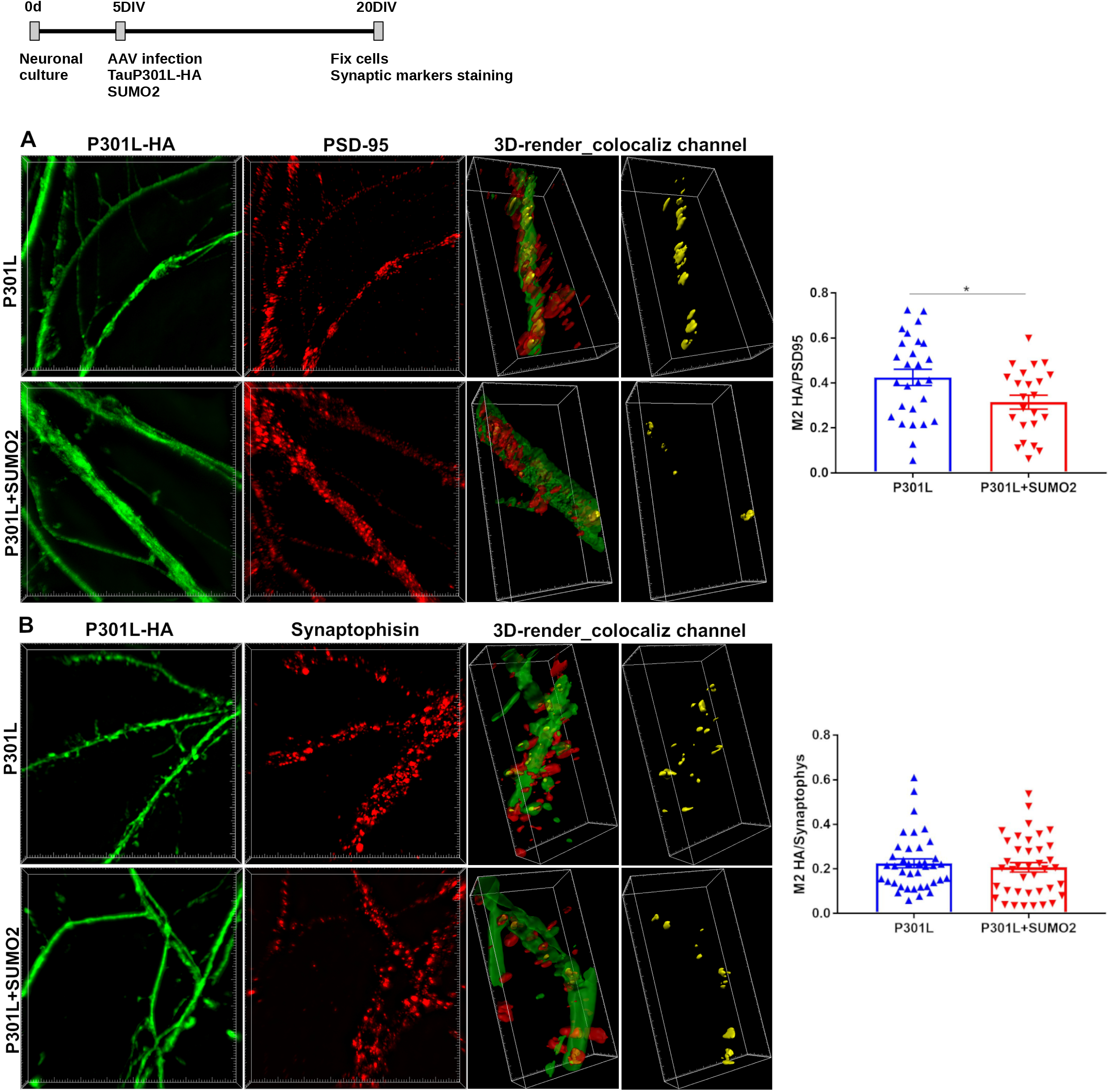
SUMO2 restores the synaptic distribution of TauP301L in neurons. Primary hippocampal neurons were transduced with 2N4R TauP301L-HA tag alone or in combination with SUMO2 AAV as in Fig 3. Representative super-resolution images of neurons stained with HA (green) antibody to detect P301L-Tau HA and PSD95 (red in **A**) or synaptophysin (red in **B**) to mark the post-synaptic or presynaptic compartment respectively. Colocalization analysis with Jacop ImageJ plugin was used to calculate Mander’s coefficient (M2). M2 is reduced for PSD95 but not synaptophysin in TauP301L+SUMO2 compared to TauP301L alone transduced neurons. The graph shows the quantification of 27 branches/condition for PSD95 (A) and 39 for synaptophysin (B) colocalizations from two independent experiments. Data are expressed as mean± SEM., unpaired t-test,*p<0.05.

### Overexpression of SUMO2 reduces the aggregation of mutant Tau in PS19 mice

Our cellular data, on the beneficial effects of SUMO2 overexpression toward mut-Tau aggregation and toxicity, prompted us to test the effects of increased SUMO2 conjugation on PS19 mice, which express the Tau P301S mutation under the mouse prion protein (*Prnp)* promoter. These mice show progressive synaptic loss followed by reduced long-term potentiation (LTP) and accumulation of Tau inclusions ^35^.

To assess the effect of SUMO2 overexpression on mutant Tau aggregation and toxicity, the PS19 mice were crossed with transgenic mice overexpressing SUMO2. SUMO2 transgenic mice are healthy, show no behavioral alterations nor lifespan changes and were characterized in Dr. Fraser’s laboratory (unpublished). Consistent with the literature, we found that overexpression of SUMO2, in the absence of any stress, did not result in increased SUMO2 conjugation *in vivo* (Supplemental Fig. 5A). Interestingly, SUMO2 conjugation, which we found reduced in PS19 mice, was restored to control levels when PS19 mice were crossed with SUMO2 overexpressing mice (Supplemental Fig. 5A). Next, we performed detergent insolubility assay in cortical and hippocampal brain extracts derived from PS19 mice and PS19/SUMO2 mice. Consistent with previous reports ^35^ and our findings obtained in cells (Fig. 3B) we found that a significant amount of P301S Tau was insoluble in sarkosyl (Fig. 5A). Moreover, we found that overexpression of SUMO2 significantly reduced the amount of phosphorylated mutant Tau in the insoluble fraction (Fig. 5A).

**Fig. 5.**
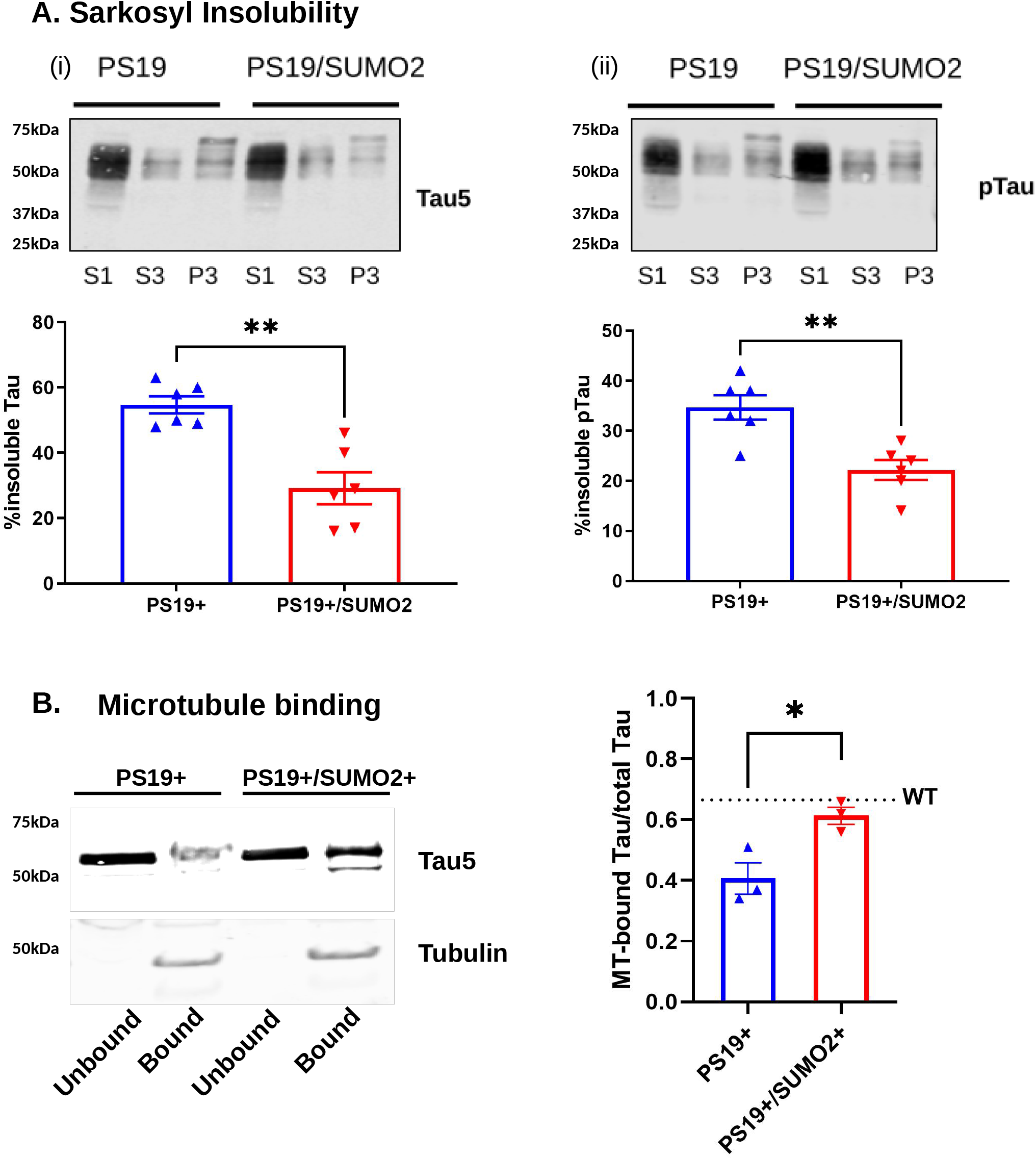
Overexpression of SUMO2 improves pathological alterations of mutant Tau in PS19 mice. **A)** (i) Representative western blot of sarkosyl soluble (S3) and insoluble (P3) fractions from mutant Tau (P301S) in PS19 and PS19/SUMO2 mice visualized with the Tau5 antibody. S1 fraction represent cleared starting homogenate. Quantification demonstrates a reduction of insoluble Tau in mice overexpressing SUMO2. (ii) Representative western blot of phosphorylated Tau (Ser396) in soluble and insoluble fractions. Quantification demonstrates a reduction of phosphorylated Tau in the sarkosyl insoluble fraction from PS19/SUMO2 mice. N=6 mice per genotype. Data in the graph are represented as fraction of sarkosyl insoluble Tau over total amount of Tau (P/S+P) and expressed as mean ± SEM, unpaired t-test,**p<0.001. **B)** Representative western blot image of microtubule (MT) binding assay performed on PS19 and PS19/SUMO2 mice. Compared to endogenous wild type Tau, (shown as dotted line on the graph) mutant Tau is significantly less associated with microtubules. Quantification demonstrates an increase of the percentage of Tau in the MT fraction in SUMO2 overexpressing mice. Tau was visualized with the Tau5 antibody. MT were visualized with Tubulin. The graph shows the quantification of three samples/group. Data are reported as mean± SEM, unpaired t-test n=3, *p<0.05.

Given that phosphorylation and aggregation of Tau are associated with reduced microtubules affinity, we performed microtubule fractionation experiments to measure the amount of P301S Tau associated with microtubules. We extracted proteins in conditions that preserve microtubules ^58^ from the cortex of Non-Tg, PS19 and PS19/SUMO2 mice. As expected, we observed a reduced percentage of mut-Tau associated with the microtubule fraction (polymerized tubulin) compared to wild-type Tau (Fig. 5B). However, we observed a significant shift in the amount of Tau toward the microtubule fraction when we measured mut-Tau in the microtubule fraction of PS19/SUMO2 mice. These data suggest that elevated levels of SUMO2 improves mut-Tau ability to bind to microtubules (Fig. 5B).

### Overexpression of SUMO2 preserves synapses in PS19 mice

Abnormally phosphorylated and aggregated Tau is toxic to synapses ^59 60^. In fact, PS19 mice are characterized by synaptic loss and synaptic plasticity impairment ^35^. Overexpression of SUMO2 significantly reduced the amount of phosphorylated and sarkosyl insoluble Tau. Thus, we asked whether SUMO2 could prevent the synaptic loss observed in the PS19 mouse model. We quantified the amount of synaptic markers that were previously reported to be reduced in PS19 mice (synapsin, PSD95 and SynGAP ^61^), using total hippocampal extracts from Non-Tg, SUMO2, PS19 and PS19/SUMO2 mice. SUMO2 mice had similar levels of synaptic proteins as Non-Tg mice, suggesting that the increased availability of SUMO2 *per se’* does not generally affect synaptic proteins because SUMO2 is not conjugated in the absence of stress (Fig. 6A and Supplemental Fig.5B). As expected, we observed a significant reduction of several synaptic markers, such as GluA2, synapsin, alpha-synuclein, synaptophysin, and SynGAP (Fig. 6A) in PS19 mice compared to Non-Tg mice. Interestingly, we found that the overexpression of SUMO2 rescued the levels of these synaptic markers in PS19/SUMO2 mice. However, SUMO2 overexpression did not rescue the reduction of PSD95 (Supplemental Fig. 5B). Overall, the changes observed were not due to neuronal loss, as the neuronal microtubule associated protein 2 (MAP2) was not reduced in PS19 mice at this time point (Supplemental Fig. 5B).

**Fig. 6.**
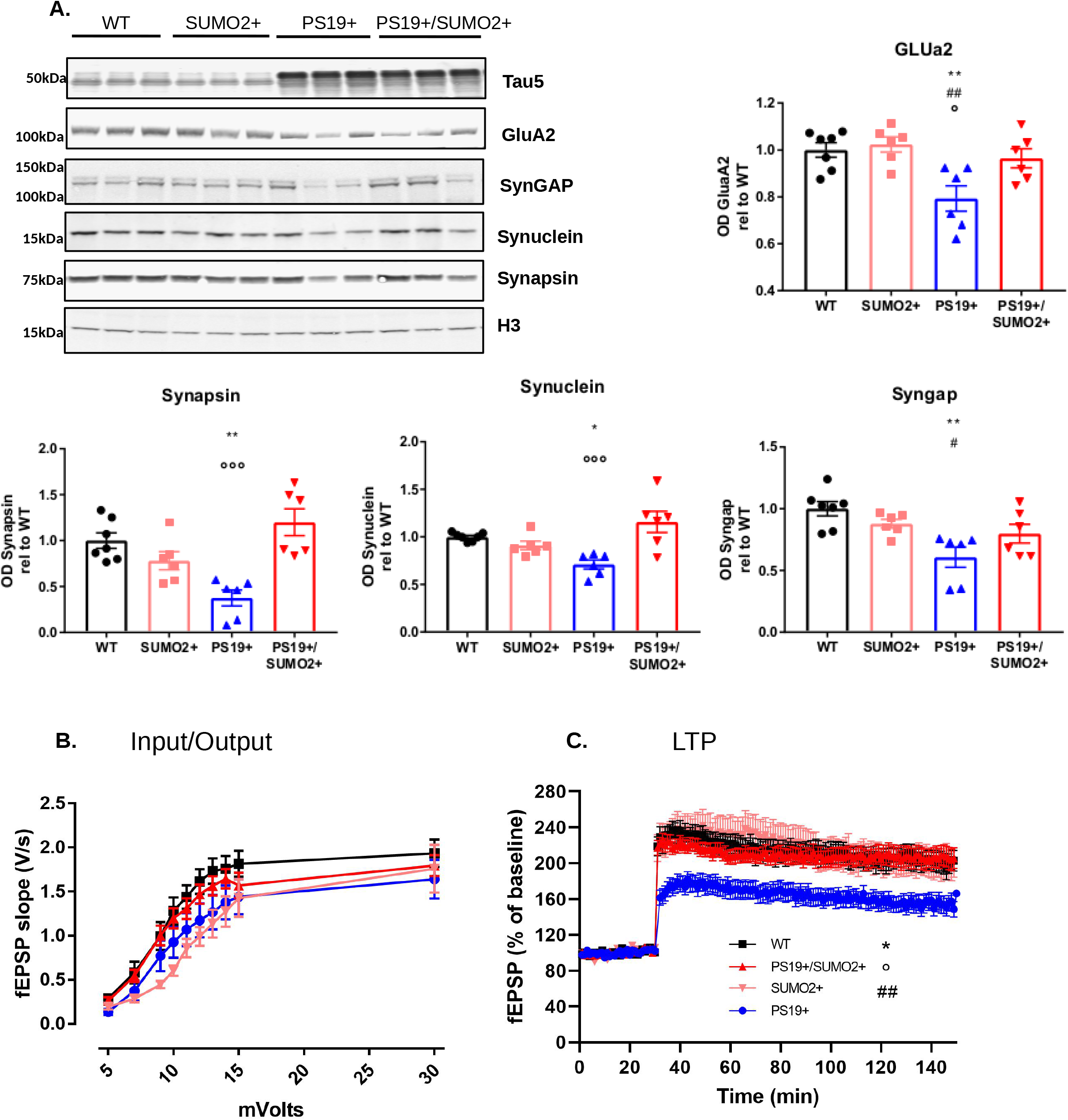
Overexpression of SUMO2 preserves synaptic markers and synaptic plasticity in PS19 mice. **A)** Representative western blot of synaptic proteins levels in 8-9 months old WT, SUMO2, PS19 and PS19/SUMO2 mice. Total hippocampal extracts were probed with antibodies against Tau, GluA2, synGAP, synapsin, alpha synuclein, and H3, as loading control. Graphs show the quantification of each synaptic marker, n=6-7 animals per genotype. PS19 mice show significant reduction of all synaptic proteins analyzed compared to WT mice. SUMO2 overexpression restores the levels of synaptic proteins in PS19 mice as demonstrated in double PS19/SUMO2 Tg mice. One-way ANOVA, followed by Tukey’s multiple comparisons post hoc test, * p< 0.05, ** p<0.01, ***p<0.001 vs Wt; ##p< 0.01 and ###p<0.001 vs SUMO2; ° p<0.05, °°p<0.01 and °p<0.001 vs PS19/SUMO2. **B)** Input-output relationship plots of hippocampal slices from WT (7-9 months old), PS19, SUMO2 and PS19/SUMO2. WT n = 12 slices (4 male, 3 female mice), PS19 n= 18 (4male, 5 female mice), SUMO2 n = 14 slices (4 male, 3 female mice), PS19/SUMO2 n=18 (5 male, 3 female mice); two-way ANOVA for repeated measures shows no differences among genotypes. **C)** LTP of hippocampal slices from WT, PS19, SUMO2 and PS19/SUMO2. The graph shows the quantification of n= 12 WT, n= 16 PS19, n= 16 SUMO2 and n= 18 PS19/SUMO2 slices. Data are the mean± SEM, two-way ANOVA for repeated measures followed by Tukey’s multiple comparisons post hoc test to compare main genotype effects, PS19 genotype *p < 0.05 vs WT ##p<0.01 vs SUMO2 and °p < 0.05 vs PS19/SUMO2 genotype.

### Overexpression of SUMO2 restores synaptic plasticity in PS19 mice

Changes in the amount of synaptic proteins are likely to underlie changes in synaptic function. Thus, we investigated the effect of mutant Tau in synaptic transmission and plasticity by analyzing the Schaffer collateral pathway in hippocampal slices from 8-9 months old mice. Basal synaptic transmission (BST) was determined by measuring the input-output relationship on the slope of the field excitatory post-synaptic potential (fEPSP) elicited through stimuli of increasing intensity. We found no difference in BST between SUMO2 and WT mice (Fig. 6B). LTP, a long-lasting form of synaptic plasticity that is thought to be associated with learning and memory ^62^, instead was markedly impaired in PS19 mice as compared to Non-Tg littermate controls (Fig. 6C). Overexpression of SUMO2 completely rescued the deficits observed in the PS19 mice (Fig. 6C).

### Overexpression of SUMO2 rescues memory impairment in PS19 mice

Since LTP experiments indicated that up-regulation of SUMO2 conjugation rescues mutant Tau-induced impairment of LTP, we aimed at extrapolating these findings to memory by assessing spatial learning and memory through two independent tasks, the object location task (OLT) ^63 64^ and the radial-arm water maze (RAWM) task. First we performed open field experiments to assess anxiety and locomotor behavior in our mice at 8-9 months of age. We did not observe any impairment, thus allowing us to further test their memory abilities (Supplemental Fig. 6A).

We evaluated the PS19 mice in the OLT with a 1-hour and 24 hr inter-trial interval. During the training phase, PS19, SUMO2, PS19/SUMO2 and Non-Tg littermates had similar total exploration times and explored the two objects equally (Supplemental Fig. 6B). Twenty-four hours after training, the animals were placed back in the arena where one of the objects was moved to a different position. Mice of all genotypes displayed similar total exploration time (Supplemental Fig. 7B). However, PS19 animals spent significantly less time exploring the object placed in a different position compared to the other groups, suggesting that the PS19 animals had impaired memory for the spatial configuration of the objects (Fig. 7A). Interestingly, the PS19/SUMO2 mice demonstrated a preference for the differently positioned object, like control mice (Fig. 7A).

**Fig. 7.**
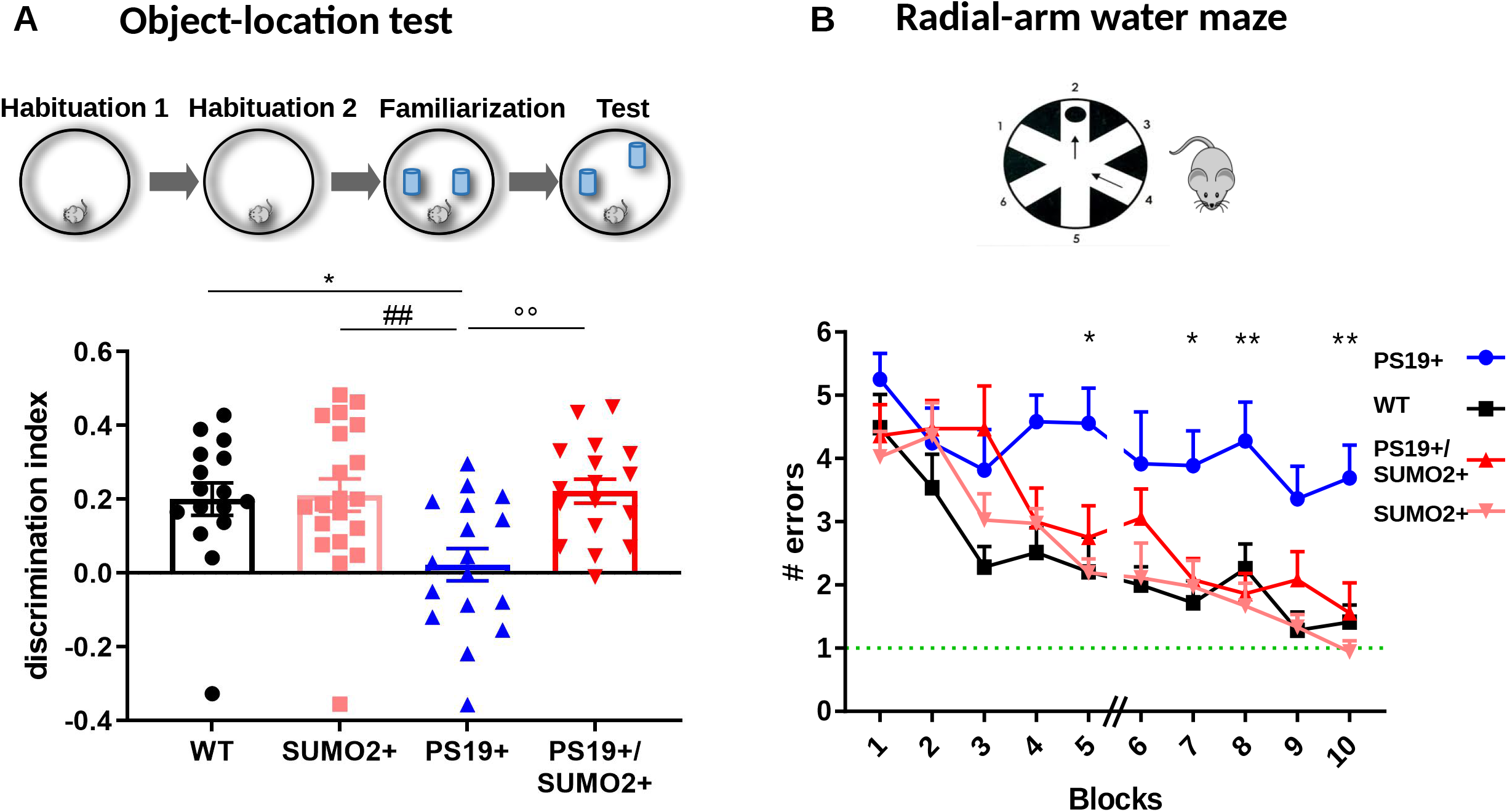
Overexpression of SUMO2 rescues memory impairment in PS19 mice. **A)** OLT: the Fig. reports the discrimination index of WT (7-9 months old), PS19, SUMO2 and PS19/SUMO2 mice. Data show that PS19 mice are severely impaired and that overexpression of SUMO2 restores memory loss. Bars represent mean ± SEM. of 16-20 mice represented by individual points. Oneway ANOVA followed by Tukey’s multiple comparisons test, *p<0.05 vs WT, ## p<0.01 vs SUMO2 and °°p<0.01 vs PS19/SUMO2. **B)** Radial arm water maze cognitive task. Number of errors by WT (7-9 months old), PS19, SUMO2 and PS19xSUMO2 mice (n=12/ genotype) to locate the hidden escape platform. Data are the mean ± SEM. of 12. mice; two-way ANOVA for repeated measures followed by Tukey’s multiple comparisons post hoc test, *p<0.05 and **p<0.01 PS19 vs PS19/SUMO2.

Next, we performed a short-term reference memory task, the RAWM ^65 66^. PS19, SUMO2, PS19/SUMO2 and Non-Tg littermates were trained to locate a platform hidden under opaque water in one of the six arms using navigational cues set in the room (Fig. 7B). Non-Tg animals made one error on average prior to successfully locating the escape platform, whereas PS19 animals made three/four errors, indicating their reduced learning capacity (Fig. 7B). Control experiments with the visible platform test excluded that PS19 have visual, motor and motivational impairment, since they showed similar swimming speed or time to find the visible platform compared to Non-Tg mice (Supplemental Fig. 6C). Overexpression of SUMO2 completely rescued the learning and memory deficits observed in the PS19 mice (Fig. 7B).

### Pharmacological treatment to increase SUMO2 conjugation rescues synaptic markers in iPSC-derived neurons expressing mutant Tau

To assess the clinical relevance of our findings, we turned to iPSC-derived neurons from patients with R406W and P301L mutations to: 1) assess the effects of mutant Tau expression on global SUMOylation; 2) determine whether a pharmacological approach could restore SUMO2/3 conjugation; and 3) test whether increased SUMO2/3 conjugation could improve mutant Tau-mediated toxicity.

First we compared the profile of SUMO conjugation in iPSC-derived neurons from patients carrying the Tau R406W mutation ^67^, and isogenic controls. We assessed both SUMO1 and SUMO2/3 conjugation in fully differentiated cortical neurons ^68^ We found a significant change in both SUMO1 and SUMO2/3 global conjugation profile, with SUMO1 being enhanced and SUMO2/3 decreased (Fig. 8A). We found similar results with the P301L mutation (Supplemental Fig. 7A).

**Fig. 8.**
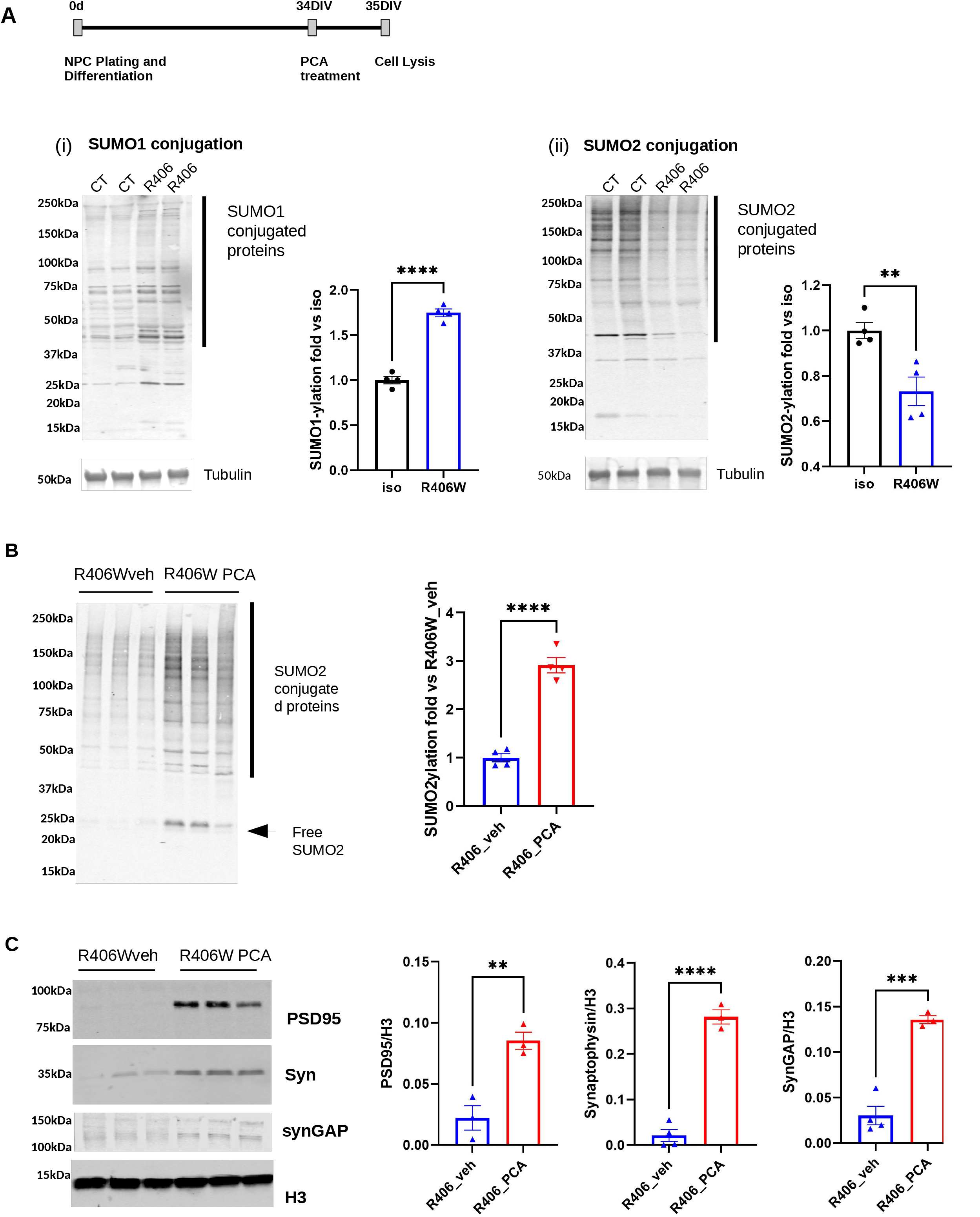
Pharmacological treatment to increase SUMOylation restores synaptic markers in iPSC-derived neurons. **A)** Representative western blots of global SUMO1 (i) and SUMO2 (ii) conjugation in iPSC-derived neurons from FTD patient with R406W tau mutation and isogenic control. Cellular extracts were probed with SUMO1 or SUMO2 specific antibodies. R406W tau-expressing neurons show significant increase of SUMO1 conjugation and reduction of SUMO2 conjugation compared to their respective isogenic controls. Tubulin is shown as a protein loading control. The graph shows the quantification of 4 samples from 2 independent experiments. Data are the mean ± SEM., unpaired t-test,**p<0,01 and *p<0.05. **B)** R406W Tau iPSC-derived neurons were treated or not with PCA (2 μM) for 24 hours. Western blot and its relative quantification demonstrate that PCA increases SUMO2-conjugation in neurons. Data are presented as means ± SEM., two-way ANOVA followed by Sidak’s multiple comparisons post hoc test, *p<0.05 and ***p<0.001. **C)** Representative western blot of synaptic markers in cellular extracts from mutated and isogenic iPSC-neurons after vehicle or PCA treatment. Blot quantification shows that PSD95, synaptophysin, and synGap protein levels are higher in PCA-than vehicle-treated R406W neurons. Data are are the mean ± SEM., of 3-4 independent experiments, unpaired t-test,*p<0.05, **p<0.01 and ***p<0.001.

Next, we treated fully differentiated cortical neurons, expressing mutant Tau or isogenic controls, with molecules that inhibit SUMO-specific proteases (SENPs), critical enzymes of the SUMO cascade responsible for removing SUMOs from target proteins. Inhibition of SENPs was previously found to increase SUMO conjugation in several cellular systems ^69^. We treated cortical neurons for 24 hrs with SENP inhibitor protochatecuic acid (PCA) and found that PCA increased global SUMO2/3 conjugation (Fig. 8B).

Interestingly, as we observed in PS19 mice, the rescue of SUMO2 conjugation was also accompanied by a recovery in the levels of synaptic proteins (PSD95, Synaptophysin, and SynGAP (Fig. 8C), thus suggesting that this pharmacological treatment was able to reproduce at least some of the improvements we observed in animal models with the genetic overexpression of SUMO2.

Taken together our findings underscore the general importance of SUMOylation, and in particular, SUMO2-mediated conjugation of Tau, in the context of mutant Tau-induced pathological changes.

## Discussion

The main findings of our study are that 1) the neuroprotective SUMO2/3 conjugation is reduced in animal models of FTD as well as in iPSC-derived neurons from FTD patients, and that 2) restoration of SUMO2/3 conjugation, achieved either genetically or pharmacologically, improves pathological outcomes in the models studied. This could be due in part to the fact that Tau is a target of SUMO conjugation and when Tau is SUMOylated by SUMO2 this reduces Tau phosphorylation and aggregation. Overexpression of SUMO2 also reduces the accumulation of mutant Tau in the post-synaptic compartment. We therefore propose that SUMO2/3 conjugation reduces mutant tau toxicity acting at different levels, all converging on restoring Tau physiological distribution and function.

### Global changes of SUMOylation in models of FTD

Following the first reports in early 2000 ^43^, a growing number of proteins associated with neurodegenerative disease were found to be SUMOylated or to co-localize with SUMO variants *in vivo* ^70^. Overall, a consensus is emerging that SUMOylation is an important PTM thanks to studies on SUMOylation by SUMO1 and SUMO2 of amyloid-like proteins such as the huntingtin protein ^23; 71^, superoxidase dismutase (SOD) or TAR DNA-binding protein-43 (TDP-43) ^72^. These reports suggest a possible contribution of the SUMOylation pathway towards the formation of higher molecular assemblies ^21^. However, some studies suggest significant differences between the different SUMO isoforms ^24 39^.

How SUMO isoforms are regulated is still poorly understood, but several studies point to a regulation at transcriptional and post transcriptional levels. In cells, SUMO1 is found mostly conjugated to substrates and is less abundant in its free form. In contrast, free SUMO2/3 levels are high under normal conditions, however, they become conjugated upon various stimuli including heat shock, arsenic treatment, and DNA damage ^73 74 75^. Under these conditions, waves of widespread but protein-specific changes in SUMOylation can take place ^76 77 78^ and are associated with improved outcomes. Furthermore, several studies identified a global increase in mainly SUMO2/3 species upon ischemic challenge, including hypothermia ^20 79 80^, which appears to protect brains against focal cerebral ischemic damage ^81 82 83^. It is therefore conceivable that a similar increase in SUMO2 conjugation might be beneficial against another type of stressor, such as amyloid-induced pathology.

Interestingly in previous work we found that SUMOylation is dysregulated in AD models and patients ^33 42 84 85^. More specifically, we found that SUMO2/3 conjugation was significantly reduced. Moreover in a recent meta-analysis study, SUMO2 appeared as a critical hub gene in a transcriptomic study of the temporal lobe from AD patients ^86^.

Here we found reduced SUMO2/3 conjugation in cortical neurons derived from patients carrying Tau mutations and in FTD animal models, suggesting that SUMO2/3 conjugation is also altered in primary Tauopathies. In support of our findings, it was recently reported that SUMO mRNA levels were altered in iPSC-derived neurons expressing mutant forms of Tau ^87^. We confirmed at the protein level that iPSC-derived neurons expressing R406W and P301L Tau have less free SUMO2 and less SUMO2-conjugated proteins. Although the precise cause of these alterations is not known, and are worth further investigations, we speculated that restoring global SUMO2/3 conjugation might result in mitigating cellular stress induced by mut-Tau.

Our studies show that Tau aggregation, hyperphosphorylation and synaptic deficits are rescued by restoring SUMO2 conjugation. We hypothesize that accumulation of abnormally phosphorylated Tau causes reduction in SUMO2/3 conjugation; the decrease in SUMO2/3 conjugation further causes Tau aggregation, leading to synaptic deficits, in a feedback loop. A possible mechanism of Tau induced-SUMO2 decrease could be the selective SUMO2 mRNA degradation following stress-dependent-mediated mRNA degradation ^88^. Pathological Tau in fact can induce ER stress ^89 90 91^, which could in turn cause the degradation of SUMO2 mRNA.

Notably, global SUMO2ylation, which is dramatically upregulated by cell stress, was proposed as a protective cellular response ^20 80 82 92^. Our experiments in PS19 mice and in iPSC-derived cortical neurons demonstrate that providing more SUMO2/3 restores SUMO2/3 conjugation, rescues the level of synaptic proteins, and prevents memory deficits caused by mut-Tau.

### Different SUMOs have different effects on mut-Tau

Identifying the key substrates responsible for the complete rescue of mut-Tau-induced pathological phenotype will be crucial to designing therapeutic approaches. One such protein could be Tau itself. Wild-type Tau was described to be SUMOylated by both SUMO1 and SUMO2/3 in cell lines and in the mouse brain ^44^.

While it was reported that SUMO1 increases Tau phosphorylation and promotes Tau accumulation in transfected cells ^36^ leading to increased aggregation and toxicity, we found that SUMO2 limits the aggregation of mut-Tau and prevents its synapto-toxicity. Although more experiments are required to better understand this phenomenon, these findings are nevertheless reminiscent of the effects of different SUMO variants on amyloid precursor protein processing by secretases leading to amyloid-beta (Aβ). Several reports describe how SUMO1, SUMO2 and SUMO3 have different effects on the levels and production of the Aβ fragment. Moreover, their overexpression plays opposite roles on Aβ-induced disruption of synaptic plasticity and memory ^29 93 94^, confirming that SUMO variants may have multiple, and sometimes opposite, roles in the biochemical regulation of amyloid proteins and their functional effect ^21^.

Interestingly our data also showed that overexpression of SUMO2 re-stabilizes normal levels of SUMO1 conjugation in total hippocampal homogenates of PS19 mice, possibly by competing with SUMO1 for the same substrates. Restoring a global equilibrium between different SUMO isoforms and their conjugates might be a critical aspect of the molecular, cellular and behavioral rescue of Tau-toxicity observed in our models.

Several questions still need to be answered before we can understand how the different forms of SUMO may influence the aggregation of Tau. Of particular interest is how SUMO2, a relatively small protein with no described enzymatic activity, can prevent aggregation, while, its close variant SUMO1, may alter the protein behavior in an opposite way. An interesting aspect is that SUMO1 and SUMO2 have different surface charges, thus their conjugation to target proteins could alter their overall structure, the total charge of the protein and interaction with binding partners ^95^. In this context, binding affinity of mut-Tau to microtubules through the 4 microtubule binding repeats, where the target lysine 340 is located, could be profoundly affected by different SUMOs. Another significant difference between SUMO1 and SUMO2 is that SUMO1 usually competes with ubiquitin, while SUMO2 instead can promote ubiquitination ^14 15 96^. Different forms of SUMOs might also alters the accessibility of residues for phosphorylation, which is greatly implicated in the aggregation and toxicity of Tau.

### Synaptic role of SUMO2

In addition to the effects of SUMO2 on mut-Tau, SUMO2 could be beneficial to neurons expressing mut-Tau by acting on additional proteins, both in the nucleus and at the synapse ^97^. In fact SUMO isoforms were detected at the synapse ^98^ and several reports indicate that SUMOylation modulates synaptic transmission ^99 100 101^. Conceivably, the improved phenotype of PS19 mice overexpressing SUMO2 may also be due to a beneficial effect of SUMO2 on synaptic function. In addition, SUMO2 overexpression may affect the activity of transcription factors, RNA binding proteins and proteins involved in signal transduction. Indeed, we found changes in SUMOylation of such proteins (Fig. 1). The transcriptional factor MyeF2, for example, is implicated in myelination and its activity depends on PTMs. Restoring MyeF2 SUMOylation could improve the myelination of axons, which is affected in animal models^102^ and in FTD patients^103 104^. It will be interesting in the future to investigate whether these changes contribute to the beneficial effects of SUMO2. Future studies on these aspects will be crucial to establish the potential implications of manipulating global SUMO conjugation as a possible therapeutic approach for Tauopathies.

## EXPERIMENTAL PROCEDURE

### HT22 Cell culture

Cells were cultured in Dulbecco modified medium (DMEM) with 10% Fetal bovine serum (FBS) and 100 U/ml penicillin-streptomycin and incubated at 37°C with 5% CO_2_.

### Plasmids for Transient Expression

Tau GFP and TauP301L GFP were obtained from addgene (Plasmid #46904, Plasmid #46908). To obtain HA-tagged Tau, Tau GFP and TauP301L GFP were subcloned into pcDNA3.1+ SUMO2-CPEB3, replacing CPEB3 using the following primers: forward primer 5’-CTAGGGATCCAATGGCTGAGCCCCGCCAG-3’; reverse primer 5-CTAGCTCGAGTTAAGCGTAATCTGGAACATCGTATGGGTAAGCAGCCAAACCCTGCTTGGC-3’). GFP-SUMO1 and GFP-SUMO2 were purchased from Addgene cloned in pcDNA3.1+. Tau P301L-K340R mutant was generated by mutagenesis of the TauP301L construct with QuickChange, following the manufacturer instructions (Forward 5’-cactgccgcctcccaggacgtgtttgata-3’ Reverse 5’-tatcaaacacgtcctgggaggcggcagtg-3’).

### Primary neuronal culture

Primary neurons from WT CD1 mice (Envigo) were cultured as previously described ^105^. Briefly, hippocampi were dissected at postnatal day 1-2, sliced into 1 mm pieces, and incubated in Hanks’ balance salt solution (Invitrogen) containing 10 units/ml papain (Sigma) at 34 °C for 30 min. Trypsin inhibitor (Sigma) was added to a final concentration of 0.5 mg/ml, and the tissue was mechanically dissociated by passing through a flame-polished Pasteur pipette. Cells were plated at 1 x 10^5^ cells/cm2 on poly-L-lysine-coated (0.1 mg/ml) plates and maintained in Neurobasal basal medium (Invitrogen) supplemented with B27 (Invitrogen), penicillin/streptomycin, and glutamine at 2 mM. To reduce the number of non-neuronal cells, ARA-C (3.3 μg/ml, Sigma) was added to the medium 48 h after plating. Neurons were washed in icecold PBS and re-suspended in the protein extraction buffer (50 mM Tris-HCl [pH7.5], 50 mM KCl, and 10mM MgCl2) supplemented with 10 mM phenylmethylsulfonyl fluoride and complete protease inhibitor cocktail (Roche Applied Science) to prevent protein degradation.

### hIPSC-derived neurons

F11362.11C11 (control) and F11362.11F10 (Tau R406W) were purchased from the Neural Stem Cell Institute (Ressaulaer, NY) as Neural Progenitor Cells (NPCs) at p2 ^68^. NHCDR ND50098 (control) and ND50099 (Tau P301L) were obtained from NINDS Human Cell and Data Repository (NHCDR) at Coriell. NPCs were grown and expanded in mTesR1 media (Stem Cell Technologies) on Matrigel (Corning)-coated wells up to p5, before being grown on (1) Matrigel-coated wells or (2) poly-L-lysine-coated glass coverslips. Cells were then differentiated at DIV0 in BrainPhys Media until maturity at DIV35.

### Adeno-associated viral transduction

Neuronal cultures were transduced at DIV5 with 3.4×10^8^ Infective Units (IU)/cm^2^ of GFP-AAV1 (Vector biolabs), SUMO2-RFP-AAV1 (Vector biolabs). And tauP301L-HA-AAV1 (Produced by the AAV Vector Unit at the International Centre for Genetic Engineering and Biotechnology Trieste (http://www.icgeb.org/avu-core-facility.html), as described previously^106^ with a few modifications. Briefly, infectious recombinant AAV vector particles were generated in HEK293T cells culture in roller bottles by a cross-packaging approach whereby the vector genome was packaged into AAV capsid serotype-1. Viral stocks were obtained by PEG precipitation and CsCl_2_ gradient centrifugation^107^. The physical titer of recombinant AAVs was determined by quantifying vector genomes (vg) packaged into viral particles, by real-time PCR against a standard curve of a plasmid containing the vector genome^108^; values obtained were in the range of 1×10^13^ vg per milliliter. At DIV20 cells were fixed for 15min at RT with paraformaldehyde 4% (PFA) +sucrose 4% for immunofluorescence or harvested for biochemical analysis.

### Immunofluorescence

Fixed neuronal cultures were rinsed three times with PBS and incubated with blocking solution (5% bovine serum albumin and 0.2% Triton X-100) for 1 h at room temperature (RT). Fixed neurons were then incubated with the primary antibodies (mouse T49 1:200 or mouse HA 1:200 for tau; rabbit Tubulin 1:200, PSD95, Synaptophysin, 1:500) overnight at 4 °C. After three rinses in PBS, neurons were incubated for 1 h RT with the secondary antibodies conjugated with Alexa/DyLight dyes (488 or 650 fluorophores, Invitrogen) and then washed 3 more times with PBS. Nuclei were stained with DAPI (2 μg/mL). Labelled neurons were kept at 4 °C until images acquisition.

### Super-resolution microscopy and image analysis

Structured illumination microscopy (SIM) was performed on a Nikon SIM system with a 100 × 1.49 numerical aperture oil immersion objective, managed by NIS elements software. Cells were imaged at laser excitation of 405 for nuclei, 488 for Tubulin and 640 nm for T49 or HA staining with a 3D-SIM acquisition protocol. Fourteen-bit images sized 1024 × 1024 pixels with a single pixel of 0.030 μm (100×) were acquired in a gray level range of 0–4000 to exploit the linear range of the camera (iXon ultra DU-897U, Andor) at 14-bit and to avoid saturation. Images were acquired over 1 μm z-stack with 0.125 μm step size. Raw and reconstructed images were analyzed with the SIM check plugin of ImageJ ^55^. SIM images were converted to 8-bit for the gray level co-occurrence matrix (GLCM) analysis. After back-ground normalization, the dendrites were manually outlined over the tubulin signal and used like the region of interest (ROI). GLCM was done on the T49 or HA signal within the ROI with the ImageJ plugin ‘GLCM analysis’(using step size = 1 pixel, step direction= 0°). Entropy, angular second moment (ASM) and inverse difference moment (IDM) were measured on 60 dendrites for each condition.

### Protein Expression and Purification

Tau expression was carried out as previously reported by ^45^ with some modifications. Briefly, BL21 E. Coli were transformed with pET-29b Tau plasmids containing full-length human Tau (2N4R) and TauP301L. Overnight liquid cultures from bacterial glycerol stock were diluted 1:40 and used for large-scale amplification. Bacteria were grown in terrific broth at 37°C to an optical density of 0.7 and expression was induced by adding 1 mM IPTG for 3 h. Bacteria were pelleted by centrifugation at 6,000 g for 20 min at 4°C and were resuspended in resuspension buffer [20 mM MES, 50 mM NaCl, 0.2 mM MgCl2, 5 mM DTT, protease inhibitor cocktail (Complete, Roche) pH 6.8]. Cells were lysed by sonication on ice for four times 15 s on/30 s off and then boiled for 20 min. Cell debris and denatured proteins were pelleted by centrifugation at 100,000 g for 1 h at 4°C. and the supernatant was stored at −80°C. Supernatants were passed through Minisart syringe filters (Sartorius) with a pore size of 0.45 μm, and loaded onto a Mono S 5/50 GL cation exchange column (GE-Healthcare). Protein was eluted in a 0.05–1 M NaCl gradient containing 20 mM Pipes (pH 6.8), 1mM MgCl2, 1 mM EGTA and 2 mM DTT. Eluates were analyzed by SDS/PAGE and fractions containing Tau protein was pooled, and 5 mM DTT was added. For further purification, samples were concentrated with Amicon Ultra 0.5mL 10.000kDa (Millipore) and loaded onto a preparative-grade Superdex 200 column (Amersham Pharmacia). Elution occurred with PBS (137 mM NaCl, 3mM KCl, 10mM Na2HPO4, 2mM KH2PO4, pH 7.4) and 1 mM DTT. Monomeric Tau was pooled and concentrated, 5 mM DTT was added and was stored at −80°C. The protein was >95% pure, as assessed by SDS/PAGE.

### In vitro SUMOylation

Recombinant human 2N4R Tau, TauP301L and TauP301S, were incubated at 37C for 2 hour together with E1, E2, SUMO2 or SUMO1 (Enzo) in the presence or absence of 5mM of ATP in SUMOylation buffer (50mM Tris-HCL pH7.5, 5mM MgCl2 and 0.02 mM DTT). The reactions were left at room temperature for an additional 16 hours before Laemmli buffer (2X) was added to the samples and analyzed by western blotting.

### Immunoprecipitation

Cells were homogenized in Lysis buffer (0.5% Triton X-100; 0.5% NP-40 and 50 mM Tris-HCl (pH 7.5) with N-Ethylmaleimide (NEM), 10 mM phenylmethylsulfonyl fluoride and complete protease inhibitor cocktail (Roche Applied Science). The homogenates (300μg) were incubated 1h at 4°C with Chemotek GFP-magnetic beads. The beads were washed four times with washing buffer for 5 min at 4°C. Bound proteins were eluted with Laemmli sample buffer and analyzed by western blotting. Cortices were homogenized in Lysis buffer following the instructions of the Pierce™ Classic Magnetic IP/Co-IP Kit. Tau protein was immunoprecipitated overnight using Tau5 antibody (5 micrograms/300 micrograms of total protein in cortical extracts).

### Detergent Insolubility Assay

We used a modified procedure described by Tatzelt et al. ^48^. Cells were lysed for 30 min at 4°C in the following buffer: 0.5% Triton X-100; 0.5% NP-40; 0.5% sodium deoxycholate; and 50 mM Tris-HCl (pH 7.5). After debris was removed by centrifugation at 16,000 × g for 10 min, the supernatant was centrifuged in a TLA 55 rotor at 65,000 rpm for 40 min. Proteins in the supernatant were precipitated with methanol, and Tau in the supernatant and insoluble fractions was analyzed by western blotting.

### 2D Agarose Gel

Cells were washed in ice-cold PBS and re-suspended in the protein extraction buffer (50 mM Tris-HCl [pH7.5], 50 mM KCl, and 10mM MgCl2) supplemented with 10 mM phenylmethylsulfonyl fluoride and Complete protease inhibitor cocktail (Roche Applied Science) to prevent protein degradation. SDS-AGE gel electrophoresis of whole-cell lysates was performed as previously described ^49^. Prior to loading on the gel, lysates (~40 μg of total protein) were incubated in the sample buffer (25 mM Tris, 200 mM glycine [pH 8.3], 2% SDS, 5% glycerol, and 0.025% bromophenol blue) at room temperature (~25°C) for 7 min. To dismantle aggregates, samples were incubated at 95°C instead of the room temperature incubation.

### Global Quantitative Proteomics analysis

For global quantitative sumo profiling of fresh frozen tissue, samples from PS19 and control littermate mice, DDA (Data dependent acquisition) based proteomics was used. In brief, frozen mouse brain tissues were lysed/ homogenized by beadbeating in lysis buffer (1% SDC, 100 mM Tris-HCl pH 8.5, and protease inhibitors) and boiled for 10 min at 95°C, 1500 rpm. Protein reduction and alkylation of cysteines, was performed with 10 mM TCEP and 40 mM CAA at 45°C for 10 min followed by sonication in a water bath, cooled down to room temperature. 3 mg of protein digestion was processed for overnight by adding alpha lytic enzyme in a 1:100 ratio (μg of enzyme to μg of protein) at 37° C and 1400 rpm. Peptides were acidified by adding 1% TFA, vortexed, and desalted using a 500 mg Sep-Pak solid-phase extraction column and dried using vacuum centrifugation.

### SUMOylated peptides enrichment

Desalted Sumolyated peptides (3 mg) were resuspended in 1.4 ml of ice-cold HS IAP bind buffer (50 mM MOPS (pH 7.2), 10 mM sodium phosphate and 50 mM NaCl) and centrifuged at maximum speed for 5 min at 4°C to remove any insoluble material. Supernatants (pH ~7.5) were incubated with the washed PTMScan® HS Ubiquitin/SUMO Remnant Motif (K-ε-GG) (Cell Signaling Technology) immunoaffinity magnetic beads for 2 hours at 4°C with gentle end-over-end rotation. After centrifugation at 2000 x *g* for 1 min, beads were washed four times with ice-cold HS IAP wash buffer and three with ice-cold HPLC water. The Sumolyated peptides were eluted twice with 0.15% TFA, desalted using SDB-RP StageTip, and dried via vacuum centrifugation.

### LC-MS/MS analysis

Desalted SUMOlyated peptides were injected in an EASY-SprayTM PepMapTM RSLC C18 50cm X 75cm ID column (Thermo Scientific) connected to an Orbitrap FusionTM TribridTM (Thermo Scientific). Peptides elution and separation were achieved at a nonlinear flow rate of 250 nl/min using a gradient of 5%-30% of buffer B (0.1% (v/v) formic acid, 100% acetonitrile) for 120 minutes with a temperature of the column maintained at 50°C during the entire experiment. The Thermo Scientific Orbitrap Fusion Tribrid mass spectrometer was used for peptide tandem mass spectroscopy (MS/MS). Survey scans of peptide precursors are performed from 350 to 1900 m/z at 120K full width at half maximum (FWHM) resolution (at 200 m/z) with a 2 x 10^5^ ion count target and a maximum injection time of 50 ms. The instrument was set to run in top speed mode with 3-second cycles for the survey and the MS/MS scans. After a survey scan, MS/MS was performed on the most abundant precursors, i.e., those exhibiting a charge state from 2 to 6 of greater than 5 x 10^3^ intensity, by isolating them in the quadrupole at 1.6 Th. We used collision-induced dissociation (HCD) with 30% collision energy and detected the resulting fragments with the auto m/z Normal scan range mode in the orbitrap. The automatic gain control (AGC) target for MS/MS was set to 5 x 10^4^ and the maximum injection time was limited to 30 ms. The dynamic exclusion was set to 30 s with a 10 ppm mass tolerance around the precursor and its isotopes. Monoisotopic precursor selection was enabled.

### LC-MS/MS Data analysis

Raw mass spectrometric data were analyzed using the MaxQuant environment v.1.6.1.0 ^109^ and Andromeda for database searches ^110^ at default settings with a few modifications. The default is used for first search tolerance and main search tolerance (20 ppm and 6 ppm, respectively). MaxQuant was set up to search with the reference mouse proteome database downloaded from UniProt. MaxQuant performed the search alpha lytic digestion with up to 4 missed cleavages. Peptide, site and protein false discovery rates (FDR) were all set to 1%. The following modifications were used for protein identification and quantification: oxidation of methionine (M), acetylation of the protein N-terminus, DiGly (K) and deamination for asparagine or glutamine (NQ).

### Extraction of Sarkosyl-insoluble Tau protein

Tissues were homogenized in 10 volumes of Tris-buffered saline (TBS: 50 mM Tris/HCl (pH 7.4), 274 mM NaCl, 5 mM KCl) with 1% protease inhibitor cocktail (Roche), 1% phosphatase inhibitor cocktail (Roche) and 1 mM phenylmethylsulfonyl fluoride (PMSF). The homogenates were then centrifuged at 27,000×*g* for 20 min at 4 °C to obtain supernatant (“SUP1”) and pellet fractions. The pellet was then resuspended in five volumes of high salt/sucrose buffer (0.8 M NaCl, 10% sucrose, 10 mM Tris/HCl, (pH 7.4), 1 mM EGTA, and 1 mM PMSF) and centrifuged at 27,000×*g* for 20 min at 4 °C. The supernatants obtained from this step were collected and incubated with sarkosyl (1% final concentration; Sigma) for 1 h at 37 °C, followed by centrifugation at 150,000×*g* for 1 h at 4 °C to obtain salt and sarkosyl-extractable (“SUP3”) and sarkosyl-insoluble (“P3”) fractions. The P3 pellet was resuspended in 50 μl TE buffer (10 mM Tris/HCl (pH 8.0), 1 mM EDTA).

### Microtubule fractionation

Frozen cortex tissue was homogenized in 10x v/w (μL:mg) RAB buffer (0.1 M MES, pH 6.8, 0.5 mM MgSO4, 1 mM EGTA, 2 mM DTT, 4 M Glycerol, 2 mM GTP, 0.1% Triton X-100). 50 mg homogenate was ultracentrifuged at 50,000 rpm at 24°C for 40 min. Supernatant (S, unbound) fraction was removed to a new microcentrifuge tube and stored at −80°C. Pellet (P, microtubule-bound) fractions were re-suspended in urea buffer (30 mM Tris-HCl, pH 8.4, 8 M urea, 1% CHAPS). 10 μL of each the supernatant and pellet fractions were analyzed by SDS-PAGE.

### Antibodies and Reagents

We used the following antibodies: HA (mouse monoclonal from Covance), GFP and Tubulin (Santa Cruz), SUMO2 (Abcam) and custom SUMO2 was a gift of Dr. Fraser, Tau (Dako), PSD95 (millipore), vinculin (millipore), P-Tau AT8 (Anaspec), GluA2, MAP2, Synaptophysin, synapsin, NR2, a-synuclein, actin, tubulin, H3. All horseradish peroxidase-conjugated secondary antibodies were from SIGMA and Santa Cruz. Alexa 488, 568 and 647-conjuated secondary antibodies were purchased from Invitrogen.

### Western Blotting

30 μg of homogenates were separated by SDS-PAGE and transferred to Immobilon-P Transfer Membrane (Millipore). Membranes were blocked in milk solution (5% milk in PBS and 0.1% tween 20). All the HRP-conjugated secondary antibodies were used at 1:5000 dilutions. Immunoblots were visualized with either Supersignal West Femto (Pierce) or Western Lightning Plus ECL (PerkinElmer). Image quantification analysis (ImageJ) was used for the calculation of protein level differences. Prism software (SAS institute) was used for statistical analysis and graph design.

### Animals

Procedures involving animals were conducted in conformity with the institutional guidelines at the Istituto di Ricerche Farmacologiche Mario Negri IRCCS, in compliance with national (D.lgs 26/2014; Authorization n. 19/2008-A issued March 6, 2008 by Ministry of Health) and international laws and policies (EEC Council Directive 2010/63/UE; the NIH Guide for the Care and Use of Laboratory Animals, 2011 edition). They were reviewed and approved by the Mario Negri Institute Animal Care and Use Committee, which includes ad hoc members for ethical issues, and by the Italian Ministry of Health (Decreto no. 420/2017-PR). Animal facilities meet international standards and are regularly checked by a certified veterinarian who is responsible for health monitoring, animal welfare supervision, experimental protocols, and review of procedures. The behavioral and electrophysiological experiments were performed at Columbia University under IACUC protocol AC-AABN7568. Breeding of the mice occurred at University of Toronto under IACUC protocol AUP6348.5 which was approved by the Animal Care Committee of the University Health Network (UHN).

### Electrophysiological recordings

Mice were sacrificed through cervical dislocation and hippocampus was removed immediately after decapitation. Transverse hippocampal slices (400 μm) were cut on a tissue chopper and transferred to the recording chamber where the physiological conditions in the brain were maintained by perfusion of ACSF continuously bubbled with 95% O_2_ and 5% CO_2_. The ACSF consisted of (in mM): NaCl (124.0), KCl (4.4), Na_2_HPO_4_ (1.0), NaHCO_3_ (25.0), CaCl_2_ (2.0), MgCl_2_ (2.0), and glucose (10.0). Slices were allowed to recover for at least 90 min before commencing the extracellular field recordings. A bipolar tungsten electrode and a glass electrode filled with ACSF were placed in the Schaeffer collateral fibers and the CA1 *stratum radiatum,* respectively. An input-output analysis was utilized to determine the maximal slope and the baseline was recorded every minute at approximately 35% of the maximum evoked slope ^<u>111</u>^. After establishing a 30 min stable baseline, LTP was induced using a theta-burst stimulation (four pulses at 100 Hz, with the bursts repeated at 5 Hz and each tetanus including three 10-burst trains separated by 15 s) and was recorded for 2 h after tetanization. LTP was measured as field-EPSP (fEPSP) slope expressed as a percentage of the baseline and the results were represented as mean±SEM. For all the electrophysiological experiments, oTau and the various drugs were diluted in the ACSF and provided through the bath solution.

### Behavioral studies

#### Radial Arm Water Maze

The RAWM consists of a white circular pool, 120 cm in diameter, filled with non-toxic white paint to make the water opaque. Within the pool there is an apparatus consisting of six arms radiating from the central area, forming six arms. Spatial cues were present on the walls of the room. Throughout the test, the water temperature was maintained stable at 24 ± 2 °C. The platform was positioned at the end of one of the arms, submerged in the water. The location of the platform (10 cm diameter) was kept constant for each mouse, while the starting position differed between the trials. The test took place for two consecutive days, and each mouse underwent 15 trials per day. On the first day, mice were trained for 15 trials, with the first 12 trials alternating between visible (platform flagged) and hidden (platform 1 cm beneath the water surface). The last 3 trials of the first day and all the 15 trials of the second day were done with hidden platform. In each trial, the mouse was allowed to swim freely for 60 s in the maze to find the platform. Once on the platform, the mouse was allowed to rest for 20 s and to observe the visual cues. If a mouse was unable to find the platform within 60 s, the experimenter guides it towards the platform for the 20 s stay. During the 1 min trial, each time the mouse entered an arm other than the goal arm (in which the platform was located), or if the mouse did not make a decision regarding which arm to explore within 10 s, an error was registered. Entry into an arm was defined as the entry of all the four paws of the mouse into the particular arm. After completing each trial, the mouse was removed from the pool, gently towel dried and placed back into its cage under a heat lamp. To avoid the learning limitations imposed by over practice and to prevent fatigue that may result from consecutive trials spaced practice training was established by running the mice in cohorts of 4 or 5 and alternating different cohorts through the 15 trials over the testing period each day. The result is shown by dividing the 30 trials into 10 blocks. Each block represents the error average of 3 consecutive trials.

#### Visible platform testing

This test was utilized for the assessment of visual and/ or motor, and/or motivational deficits. It was performed in the same pool as the RAWM. The test takes place in 2 consecutive days and mice underwent 2 sets of trials each day. Every set consisted of 3 trials in which the mouse trained to find the visible escaping platform, flagged with a bottle cup on the top. During one set of trials the platform was located in one of the three quadrants of the pool. The starting position of the mouse was kept constant for a specific location of the platform. The mice were placed gently on the water, facing the walls, and each trial lasted until the mouse had found the platform until the maximum time of 60 s. After the end of the trials mice were guided to the platform and allowed to observe environmental cues for 20 s. Time between entering the pool and reaching the platform (latency) and velocity were analyzed by using a video tracking system (Ethovision XT). The results were shown in 4 blocks and each block represents the average of one set of experiment.

#### Open field

The test has been used for assessing exploratory behavior and anxiety levels ^112^. Mice were placed in a novel open environment consisting of Plexiglass transparent walls (model ENV-520; Med Associates, St. Albans, Vermont) (43.2 cm long × 43.2 cm width × 30.5 cm high). Their activity was automatically recorded for 10 min, on two consecutive days.

#### Object Location (OLT)

The apparatus for the OLT test consists of a circular arena, 40 cm in diameter in which the back half is made of white polyvinyl chloride (PVC) and the front half is made of transparent PVC. The test was conducted in a dark room with a red fluorescent bulb for illumination. We used 2 different sets of two identical objects, which were divided in a semi-random manner between animals. The objects consisted of (1) a massive metal rectangular prism (2.5 cm × 5 cm × 7.5 cm) containing two holes (diameter 1.5 cm) and (2) a massive aluminum cube with a tapering top (4.5 cm × 4.5 cm × 8.5 cm). Prior to the testing, the animals were habituated to handling and the arena for two days. During habituation, mice were introduced to the arena for ten minutes to explore the first or the second set of objects. Afterwards, the mice were tested in the short- and the long-lasting version of the test in order to access changes in short- and long-term memory, respectively. For the short version of the test, mice were placed in the arena containing one set of identical objects for 10 minutes (Trial 1: T1) and after 1 hour inter-trial interval were placed back in the arena for the test trial (Trial 2: T2) that lasted also 10 minutes. During T1, one set of identical objects were placed in the middle of the arena, having the same predetermined distance with each other, while during T2, one of the two objects from T1 was moved to a different position on a vertical line, closer to the wall of the arena. For assessing long-term memory, mice underwent the same test, but with 24 hours interval between T1 and T2. During T1 and T2, the time that the mice spend exploring each object was manually scored. Next, we calculated the total time that the mice spent exploring the objects during T1 and T2 and the discrimination index. The discrimination index represents the difference in exploration time between the two objects during T2, divided by the total exploration of the two objects during T2.

#### Statistical analysis

For electrophysiological recordings, results were analyzed by ANOVA for repeated measures comparing traces after tetanic stimulation with treatment condition as the main effect. For the behavioral tests, animals were run in cohorts in which sex of mice was kept balanced across groups. Results were analyzed with ANOVA for repeated measures (RAWM errors and latency) or one-way ANOVA with Bonferroni post-hoc correction. For western blotting, conditions were compared by using unpaired t-test or one-way ANOVA. Statistical analysis was performed by using Systat 9 software (Chicago, IL, USA). Differences were considered significant at a *p* value less than 0.05. Results were expressed as Standard Error of the Mean (SEM).

## ACKNOWLEDGMENTS

We thank Kathy Ha and Hakyung Koo for assistance with animal husbandry; Barbara Corneo, Director of Columbia Stem Cell Core Facility, for help with differentiating TauP301L iPSC into cortical neurons; we thank Luca Colnaghi for subcloning TauP301L-HA into AAV vector (Addgene #58880) and for helpful discussion during the early stages of the project; Daniel Metzger for technical help in subcloning of the mutant Tau; Marco Bosica for technical assistance. We also thank Stefano Fumagalli for advice with Superresolution Imaging and analyses. This work was supported by the Telethon foundation (TCP11015 to L.F.), the Alzheimer Association (AARG 17-505136 to L.F.), Canadian Institutes of Health Research (PJT-173497 to P.E.F.), the Howard Hughes Medical Institute (E.R.K.), National Institute of Health (NINDS R01NS110024 to OA) (NINDS R01 NS082730 to NMK).

## AUTHOR CONTRIBUTIONS

LF conceived the study. LF, FO, EA, LKF, RS designed, performed, and analyzed experiments. ER performed experiments. HT and SM generated the SUMO2 transgenic mice. LZ purified TauP301L-HA AAV particles. NMK provided recombinant Tau (WT, P301L/S). TK provided funds. RC, GF, KK, NMK, ERK, PEF, and OA provided critical insight and helpful discussion. LF coordinated the study with input from PEF and OA. LF wrote the manuscript with input from all authors. All authors approved the final manuscript version.

