## Supplemental Figures for "SUMO2 Protects Against Tau-induced Synaptic and Cognitive Dysfunction"

### Supplemental Figure 1

A

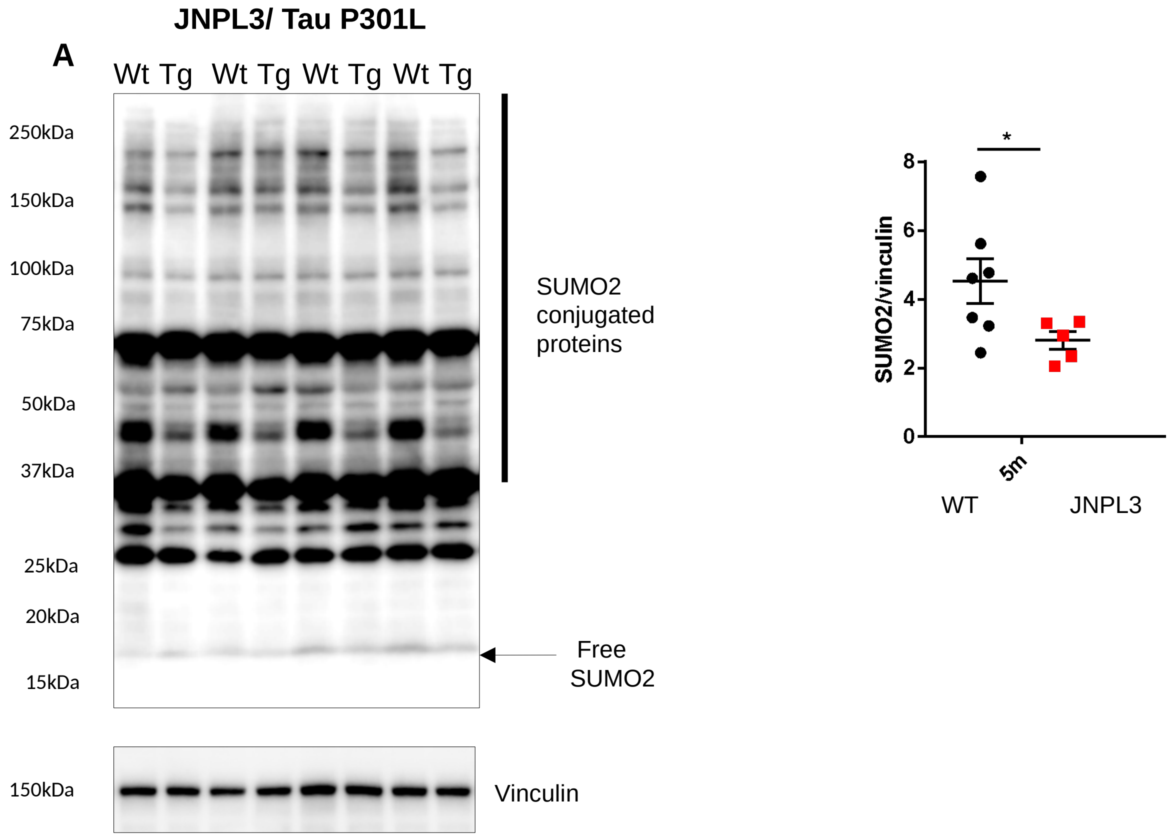


B **Mutant Tau are substrate of SUMOylation *in vitro***

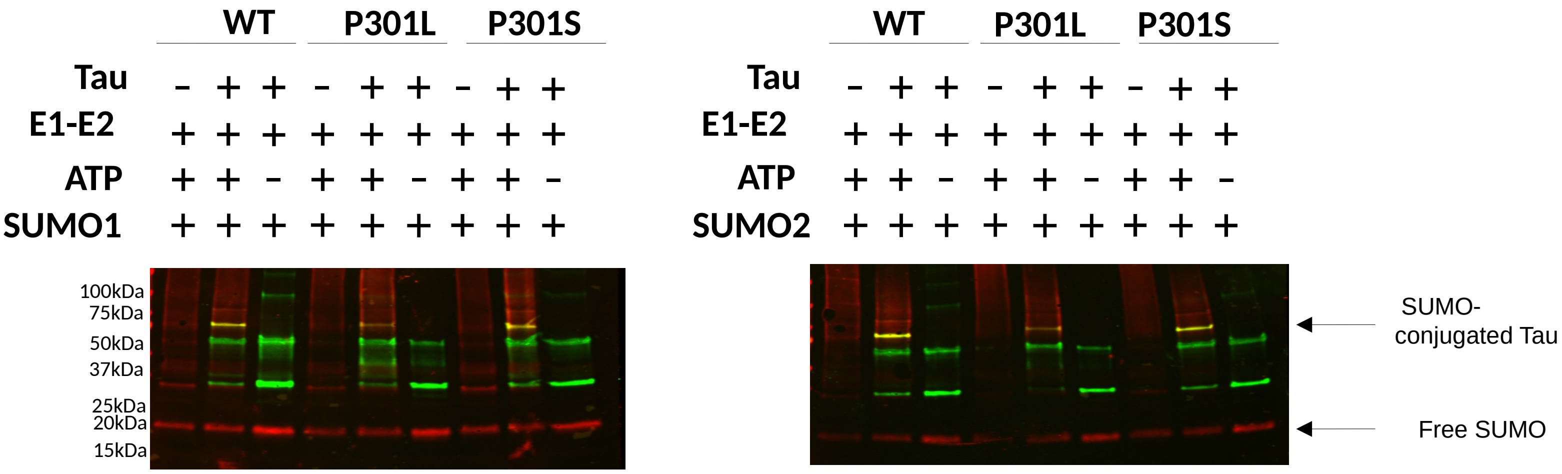

**Supplemental Figure2**

A

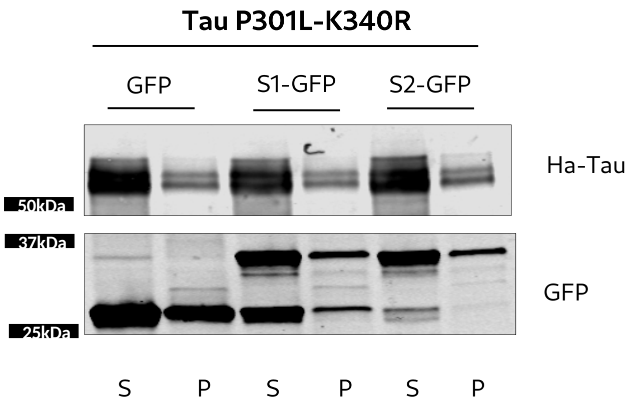

**Supplemental Figure 3**

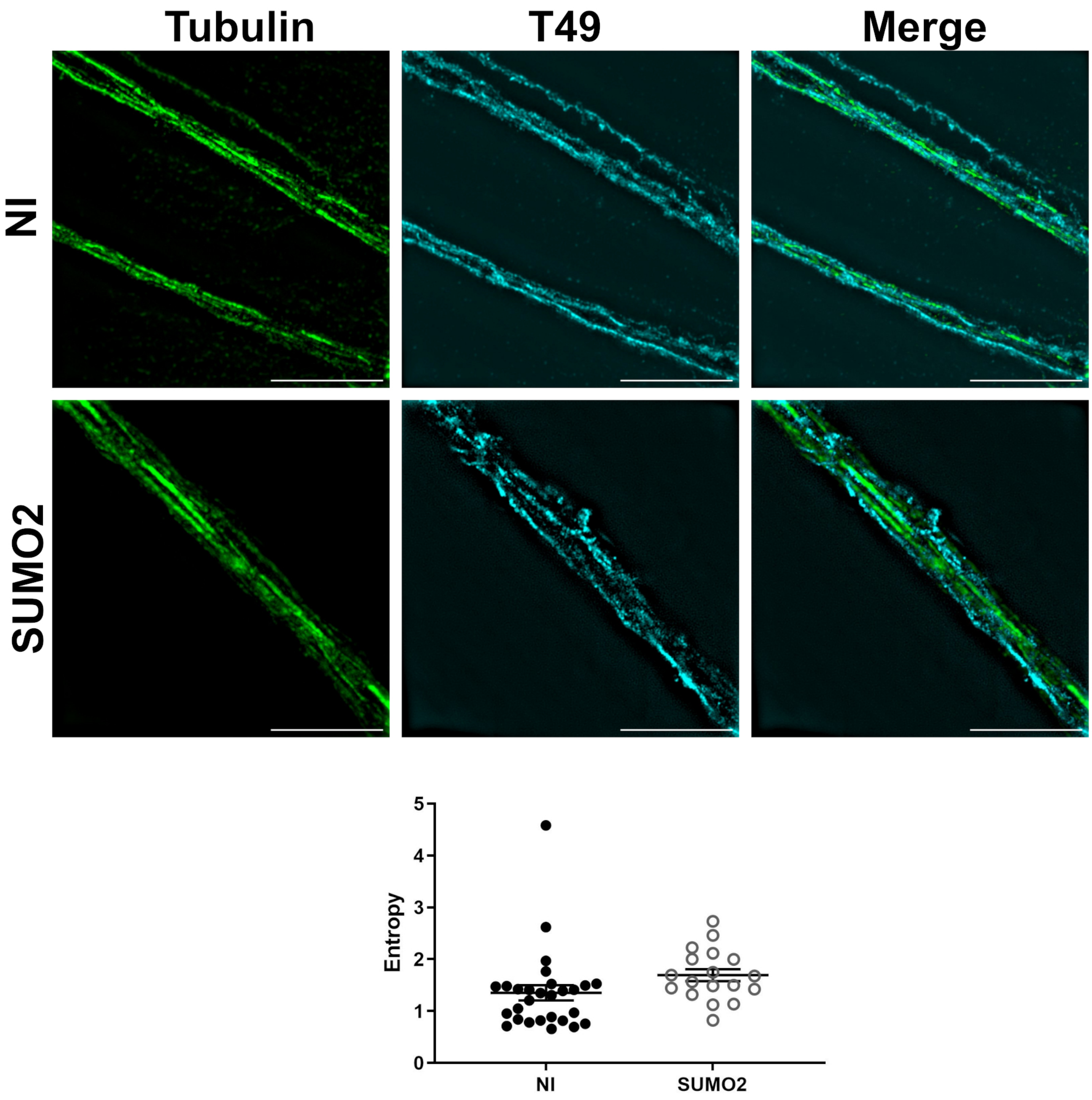

**Supplemental Figure 4**

A

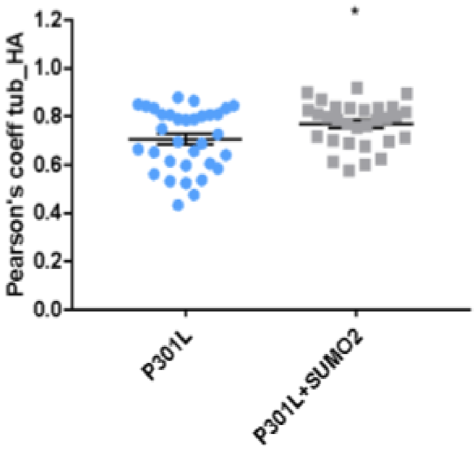

#### Supplemental Figure 5

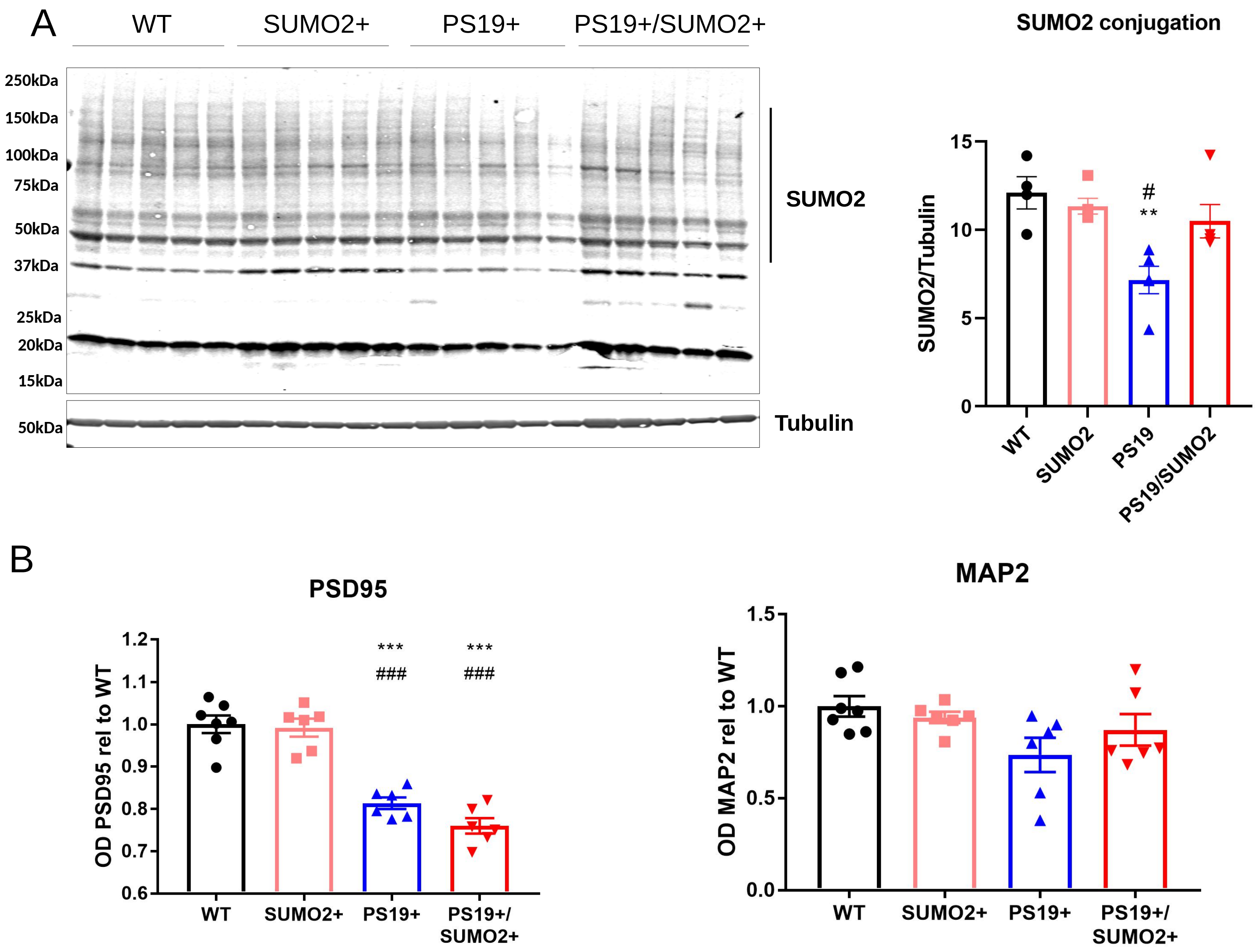

#### Supplemental Figure 6

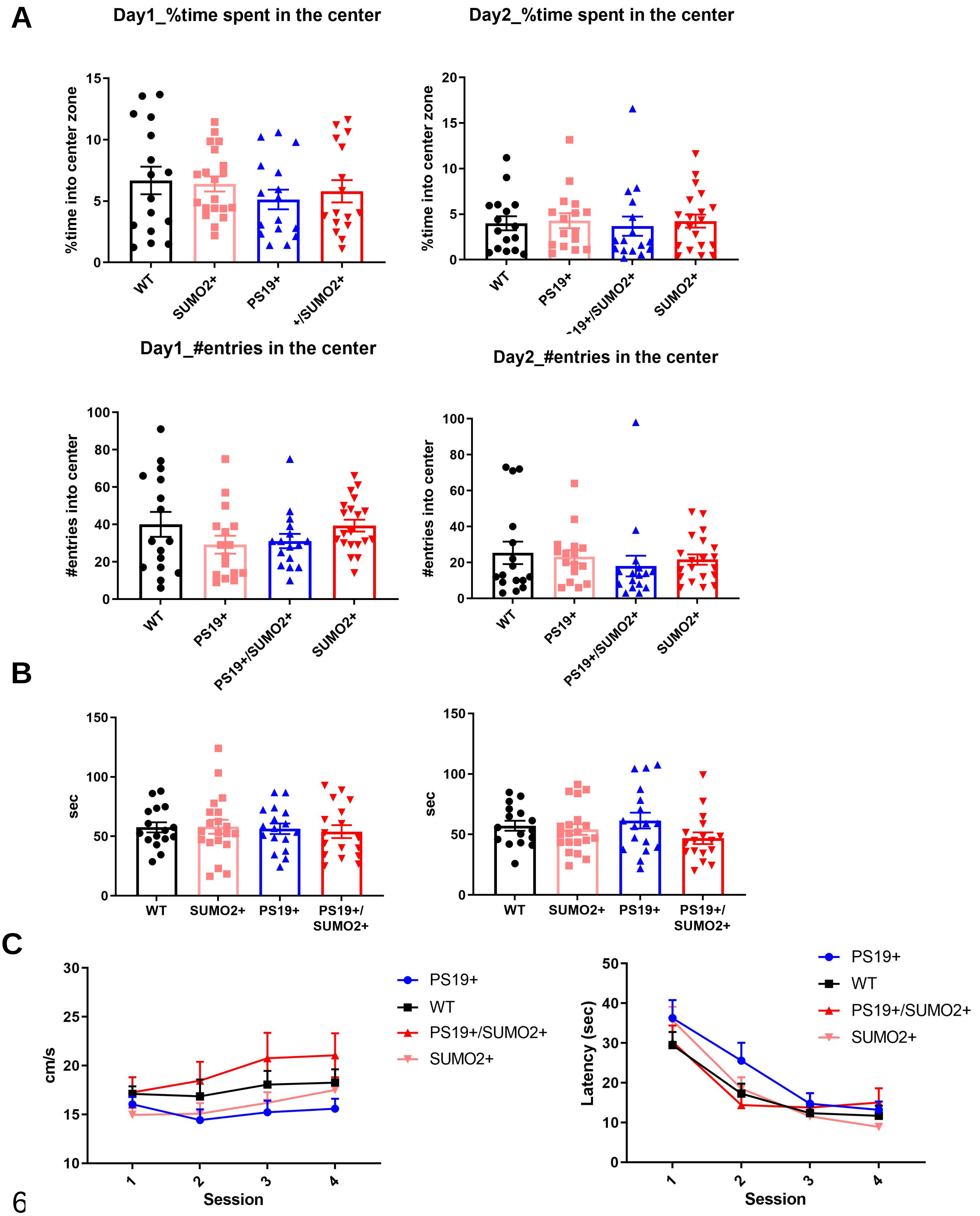

### Supplemental Figure 7

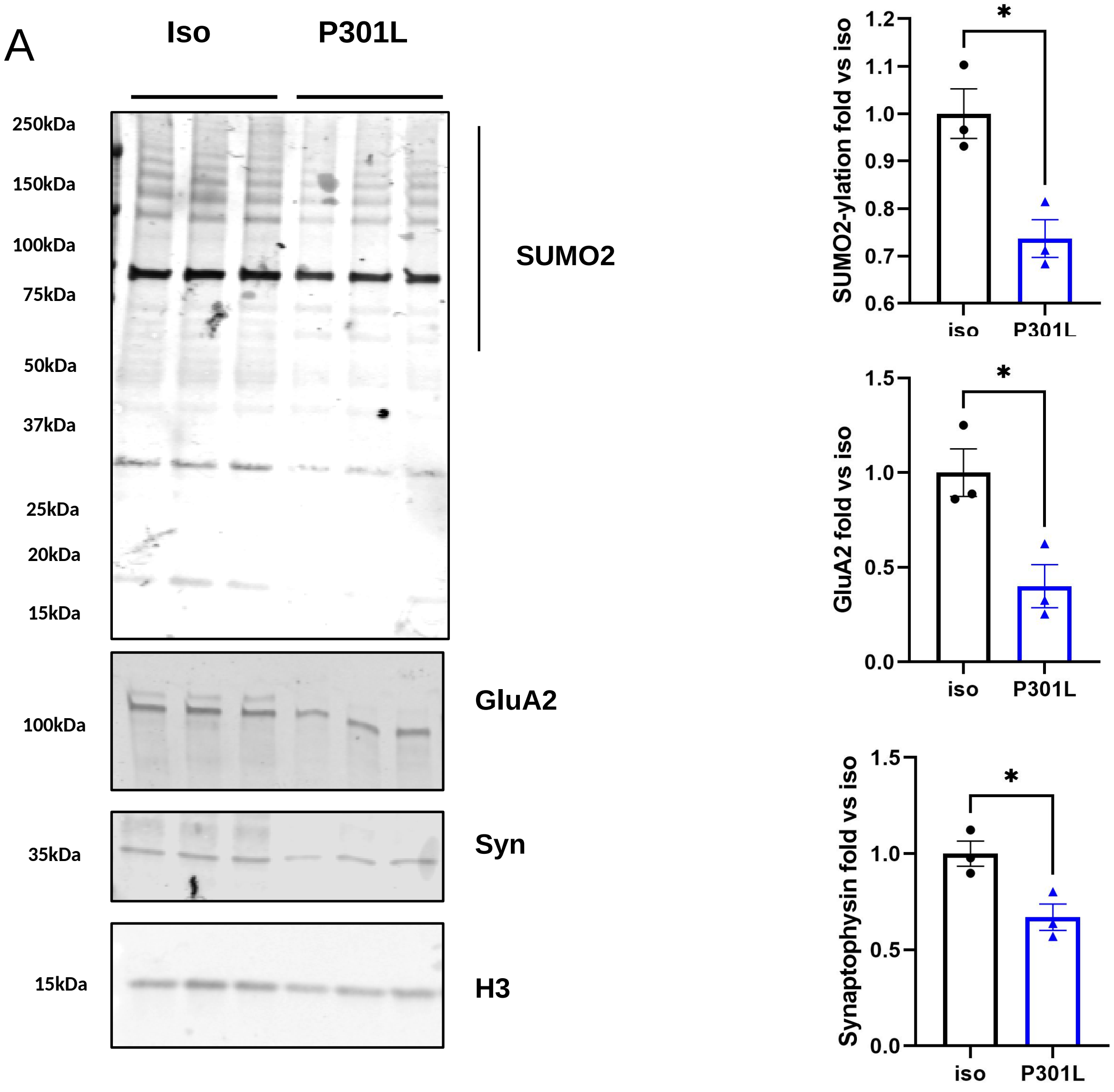
