## Supplemental material (Figures legends) for "SUMO2 Protects Against Tau-induced Synaptic and Cognitive Dysfunction"

**Supplemental Figures Legend**

**Suppl. Fig.1. Global SUMO2 conjugation is altered in JNPL3 mice. A)** Representative western blot of global SUMO2 conjugation in JNPL3 mice overexpressing the human Tau protein with the P301L mutation. Total cortical extracts were probed with SUMO2 specific antibodies. Transgenic mice show significant reduction of SUMO2 conjugation compared to their respective controls. Vinculin was used as protein loading control. Data are the mean ± SEM. of 5-7 animals per genotype; unpaired t-test, *p< 0.05. **B)** In vitro SUMO conjugation of tau proteins; WT Tau hTau40 (2N4R) as well as the corresponding proteins harboring the P301L and P301S mutations were subjected to in vitro SUMOylation combining Tau, E1-E2 enzymes, ATP and SUMO isoforms. Red signal corresponds to SUMO proteins, green correspond to Tau proteins. All Tau proteins can be modified by SUMO1 and SUMO2, as shown by the yellow band.

**Suppl. Fig. 2. SUMO2 directly counteracts aggregation of TauP301L.** Representative Western blot probed with HA antibody showing detergent insolubility of TauP301L-HA protein from cell extracts of overexpressing cells. Soluble and insoluble (pellet) fractions are labelled as 'S' and 'P' respectively. HEK293 cells were transfected with HA tagged TauP301L with K340R mutation to prevent SUMOylation, together with GFP, GFP-SUMO1 (S1-GFP) or GFP- SUMO2 (S2-GFP). SUMO2 has no effects on the solubility of TauP301L when acceptor lysine 340 is mutated**.**

**Suppl. Fig. 3. SUMO2 reduces the aggregation of TauP301L in cultured primary hippocampal neurons.** Graph shows quantification of co-occurrence of Tau P301L signal (HA) and tubulin. Pearson's coefficient was used to assess co-occurrence. TauP31L is more associated with Tubulin when neurons overexpress SUMO2. The graph shows the quantification of 33-45 branches from 2 independent experiments for TauP301L and TauP301L+SUMO2 neurons respectively. Data are expressed as mean± SEM. unpaired t-test. *p<0.05.

**Suppl. Fig. 4. SUMO2 does not alter entropy of endogenous Tau in cultured primary hippocampal neurons.**

Representative super-resolution images of endogenous Tau signal (T49 antibody, cyan) and tubulin (green) in not infected (NI) or SUMO2 AAV infected neurons. No changes in the appearance of Tau distribution were observed. Gray-level co-occurrence matrix (GLCM) was used for the quantification of tau signal distribution in dendrites. GLCM analysis showed similar entropy in NI and SUMO2 infected neurons. The graph shows the quantification of 33-45 branches for NI and SUMO2 neurons respectively from two independent experiments. Data are expressed as mean± SEM., unpaired t-test, ns.

**Suppl. Fig. 5. Overexpression of SUMO2 preserves synaptic markers and synaptic plasticity in PS19 mice. A)** Representative western blot images of SUMO2 conjugated proteins levels in

Non-Tg (WT), SUMO2, PS19 and PS19/SUMO2 mice. SUMO2 conjugation is restored in PS19 mice crossed with SUMO2. **B)** Total hippocampal extracts were probed with anti PSD95, , and MAP2antibodies. Graphs show the quantification of each synaptic marker, mean ± SEM., of 6-7 animals per genotype. One-way ANOVA, followed by followed by Tukey's multiple comparisons post hoc test, * p< 0.05, ** p<0.01, ***p<0.001 vs Wt; #p<0.05, ##p< 0.01 and ###p<0.001 vs SUMO2; ° p<0.05 vs PS19/SUMO2.

**Suppl. Fig. 6. Overexpression of SUMO2 preserves spatial memory in PS19 mice. A)** Open field analyses show that all groups of animals have normal locomotion and no anxiety. Percentage of time spent in the center and number of entries in the center over the course of two consecutive days, measured for WT, PS19, SUMO2 and PS19/SUMO2 mice. Data are the means ± s.e.m., one-way ANOVA followed by Tukey's multiple comparisons post hoc test shows no differences between groups. **B)** Total exploration in the OLT task was similar in all 4 groups. Data are the means ± SEM., one-way ANOVA followed by Tukey's multiple comparisons post hoc test shows no differences between groups. **C)** Speed and latency in the visible platform. Data are the means ± s.e.m., two-way ANOVA for repeated measures followed by Tukey's multiple comparisons post hoc test, show no differences were observed between groups.

**Suppl. Fig. 7. Pharmacological treatment to increase SUMOylation restores synaptic markers in iPSC-derived neurons. A)** Representative western blots of global SUMO2 conjugation in iPSC-derived neurons carrying the P301L tau mutation and isogenic control. P301L tau-expressing neurons show significant reduction of SUMO2 conjugation. H3 was used as a protein loading control. **B)** The graph shows the quantification of 3 samples. Data are the mean ± s.e.m., unpaired t-test, *p<0.05.  Blot quantification shows that GluA2 and synaptophysin, protein levels are reduced in P301L neurons. Data are are the mean ± SEM., of 3 independent replicates, unpaired t-test,*p<0.05.
